# Open Biomedical Network Benchmark A Python Toolkit for Benchmarking Datasets with Biomedical Networks

**DOI:** 10.1101/2023.01.10.523485

**Authors:** Renming Liu, Arjun Krishnan

## Abstract

Over the past decades, network biology has been a major driver of computational methods developed to better understand the functional roles of each gene in the human genome in their cellular context. Following the application of traditional semi-supervised and supervised machine learning (ML) techniques, the next wave of advances in network biology will come from leveraging graph neural networks (GNN). However, to test new GNN-based approaches, a systematic and comprehensive benchmarking resource that spans a diverse selection of biomedical networks and gene classification tasks is lacking. Here, we present the Open Biomedical Network Benchmark (OBNB), a collection of benchmarking datasets derived using networks from 15 sources and tasks that include predicting genes associated with a wide range of functions, traits, and diseases. The accompanying Python package, `obnb`, contains reusable modules that enable researchers to download source data from public databases or archived versions and set up ML-ready datasets that are compatible with popular GNN frameworks such as PyG and DGL. Our work lays the foundation for novel GNN applications in network biology. `obob` will also help network biologists easily set-up custom benchmarking datasets for answering new questions of interest and collaboratively engage with graph ML practitioners to enhance our understanding of the human genome. OBNB is released under the MIT license and is freely available on GitHub: https://github.com/krishnanlab/obnb

## 1 Introduction

Life is orchestrated by remarkably complex interactions between biomolecules, such as genes and their products. Network biology [6, 7, 43, 90] has demonstrated remarkable success over the past two decades in systematically uncovering genes’ functions and their relations to human traits and diseases. Accurately identifying genes associated with a particular disease, for example, is a vital step towards understanding the biological mechanisms underlying the condition, which in turn could lead to novel and effective diagnostic and treatment strategies [93, 51]. Early work in network biology focused on network diffusion-type methods based on the *guilt-by-association* principle [67], which states that genes interacting with each other likely participate in the same biological functions or pathways. The performance of these methods has subsequently been improved by the application of supervised learning [65]. In this trajectory, the next wave of improvements in network biology is likely to emanate from the surge of powerful graph machine learning (ML) techniques such as graph embeddings [27, 14] and graph neural networks [97, 102]. These methods have shown promising results in many graph-structured tasks, such as social networks [34], and have started to attract researchers to apply them to biological tasks [5, 100, 66]. To this end, accelerating the development and application of graph ML methods in network biology is of great importance.

However, there is a critical need for standardized benchmarks that allow reliable and reproducible assessment of the novel graph ML methods [78, 34]. Recent efforts such as MoleculeNet [96] and

Preprint. Under review.

Therapeutics Data Commons [37] for molecular and therapeutics property predictions, and Benchmarking GNN [23] and Open Graph Benchmark [34, 35] for more general graph benchmarks, have proven valuable in advancing the field of graph ML by providing carefully-constructed benchmarking datasets for applying specialized methods. Meanwhile, such comprehensive benchmarking datasets and systems are currently lacking for network biology. Furthermore, setting up ML-ready datasets for network biology is incredibly tedious. Some necessary steps include converting gene identifiers (IDs), setting up labeled data from annotated biomedical ontologies, filtering labeled data based on network gene coverage, and constructing realistic data splits mimicking real-world biologically meaningful scenarios. As a result, despite the remarkable amount of publicly available data for biomedical networks [36, 44] and annotations [79], the only widely available ML-ready datasets for network biology are `PPI` dataset from OhmNet [104] and `PPA` from the Open Graph Benchmark [34].

**Contributions** In this work, we address this critical need. Our main contributions are as follows:

1. We present a Python package `obnb` that provides reusable modules for data downloading, processing, and split generation to set up node classification benchmarking datasets using publicly available biomedical networks and gene annotation data. The first release version contains interfaces to networks from 15 sources and annotations from three sources.
2. We present a comprehensive benchmarking study on the OBNB node classification datasets with a wide range of graph ML approaches and tricks to set up the baseline for future comparisons.
3. We analyze the benchmarking results and point out several exciting directions and the potential need for a special class of graph neural network to tackle the OBNB tasks.

### 1.1 Related work

Several existing Python packages share similar goals with `obnb`, primarily focusing on establishing biomedical network datasets and facilitating their analyses. In these networks, nodes typically represent genes or their products, such as proteins, while edges represent the functional relationships between them [36], such as physical interactions. `PyGNA` [24] offers a suite of tools for analyzing and visualizing single or multiple gene sets using biological networks. `PyGenePlexus` [62] and `drug2ways` [76] specialize in network-based predictions of genes and drugs. The `OGB` [34] platform houses a variety of graph benchmarking datasets, which includes a PPI dataset akin to the STRING-GOBP dataset in OBNB (Table S10). Nonetheless, all the aforementioned packages have a limited number of, if any, biomedical network and label data. `obnb`, on the other hand, provides an extensive number of biomedical networks and diverse gene set collections to facilitate the systematic evaluation of graph machine learning methods on diverse datasets. In a related domain, `PyKEEN` [3] provides a vast array of biomedical knowledge graph (KG) datasets and KG embedding methods. There, the main task of interest is link prediction, through which the missing knowledge link can be completed. Other notable works for constructing large-scale biomedical KG and setting up link-prediction benchmarks from them include `BioCypher` [56] and `OpenBioLink` [11]. While the tasks associated with KG [92] are orthogonal to the node classification settings, it is possible to reformulate gene classification problems as KG completion and vice versa [5, 100]. Nevertheless, the advantages and drawbacks of these two approaches are yet to be comprehensively evaluated.

## 2 Systems description

Making the process of setting up ML-ready network biology benchmarking datasets from publicly available data as effortlessly as possible is the core mission of `obnb`. We implement and package a suite of graph (`obnb.graph`) and label (`obnb.label`) processing functionalities and couple them with the high-level data object `obnb.data` to provide a simple interface for users to download and process biological networks and label information. For example, calling `obnb.data.BioGRID(“datasets”)` and `obnb.data.DisGeNET(“datasets”)` will download, process, and save the BioGRID network and DisGeNET label data under the `datasets/` directory, which can then be loaded directly next time the functions are called. Users can compose a dataset object using the network and the label objects, along with the split (Section 2.3), which can then be used by a model trainer to train and evaluate a particular graph learning method. Alternatively, the composed dataset object can be transformed into data objects for standard GNN framework, including PyTorch Geometric (PyG) [25] and Deep Graph Library (DGL) [91].

In the following subsections, we go into details about the processing steps for the biomedical networks (Section 2.1), creation of node labels from annotated ontologies (Section 2.2), and preparation of data splits (Section 2.3). Finally, as we are dedicated to creating a valuable resource for the community, we closely follow several open-source community standards, as elaborated in Appendix A.6.

### 2.1 Network

#### Downloading

Currently, tens of genome-scale human gene interaction network databases are publicly available, each constructed and calibrated with different strategies and sources of interaction evidence [36]. Unlike in many other domains, such as chemoinformatics, where there are only a few ways to construct the graph (e.g., molecules), gene interaction networks can be defined and created in a wide range of manners, all of which capture different aspects of the functional relationships between genes. Some broad gene interaction mining strategies include experimentally captured physical protein interactions [82], gene co-expression [45], genetic interactions [19], and text-mined gene interactions [83]. We leverage the Network Data Exchange (NDEx) [75] to download the biological network data when possible (Figure 1a). The obtained `CX` stream format is then converted into a `obnb.graph` object for further processing.

**Figure 1.**
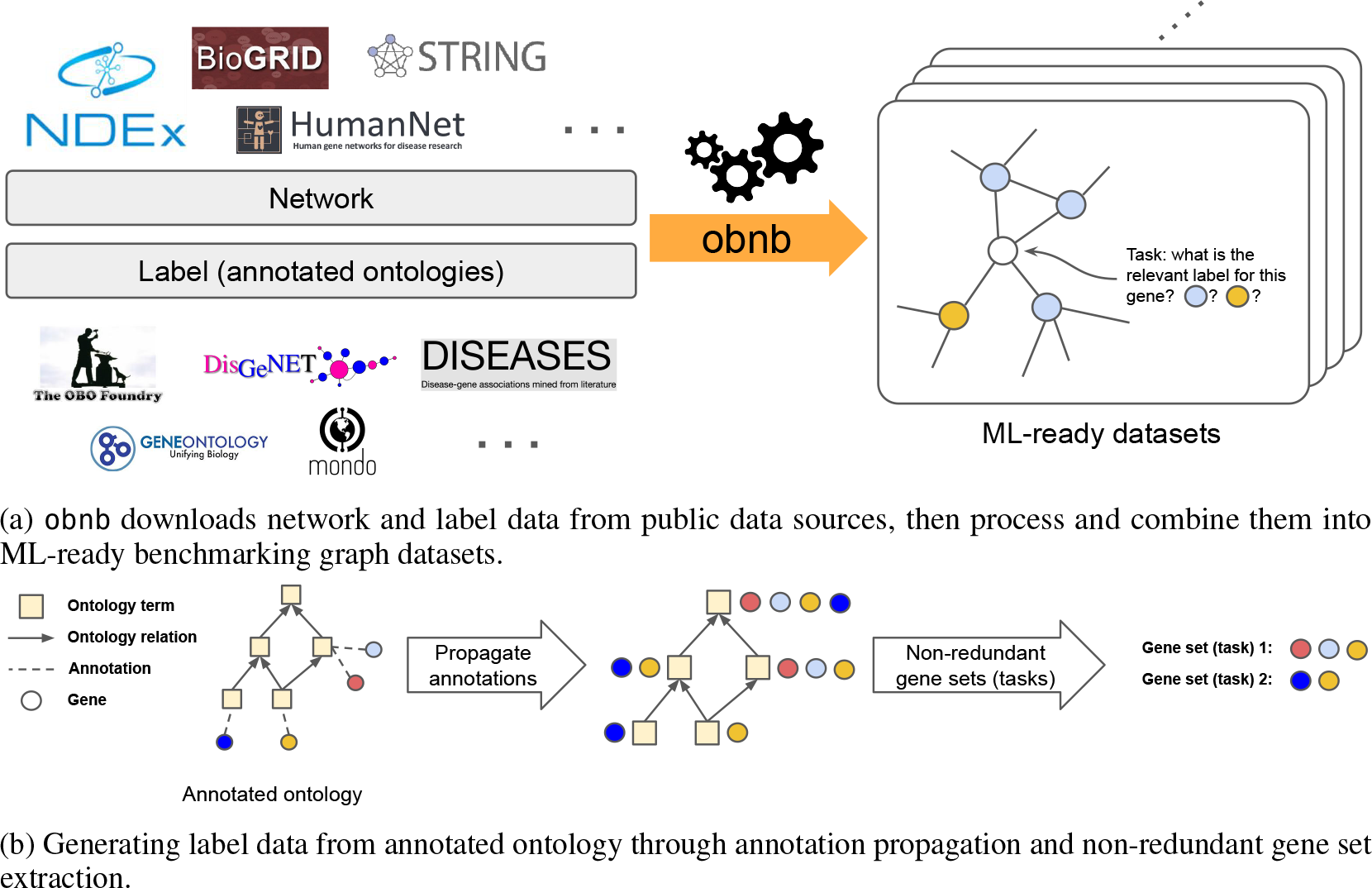
Overview of `obnb` data processing and benchmarking dataset generation

#### Gene ID conversion

In gene interaction networks, each node represents a gene or its gene product, such as a protein. There have been several standards in mapping gene identifiers, and different gene interaction networks might not use the same gene ID. For example, the STRING database [83] uses Ensembl protein ID [40], PCNet [36] uses HGNC [28], and BioGRID [82] uses Entrez gene ID [40]. Here, we use the NCBI Entrez gene ID for its advantages, such as supporting more species other than Human and being less ambiguous [33]. To convert other types of gene IDs into the Entrez gene, we use the MyGene query service [95], which provides the most up-to-date gene ID mapping across tens of gene ID types. Following, we remove any gene where more than one gene ID is mapped to the same Entrez gene, which indicates ambiguity in annotating the gene identifier currently. The gene interaction network after gene ID conversion will contain equal or less genes, all of which are one-to-one mapped from the original gene IDs to the corresponding Entrez genes.

#### Connected component

In practice, a small fraction (typically within 1%) of genes may be disconnected from the largest connected components. The disconnectedness of the network is typically due to missing information about gene interactions from the measurement; the more information a network uses to define the interactions, the denser and less likely there will be disconnected genes. Thus, we extract the largest connected component of the gene interaction network by default as it is more natural to have a single component for the transductive node classification tasks we consider. However, a user can also choose to take the full network without extracting the largest connected component by specifying the `largest_comp` option to `False` when initializing the data object.

### 2.2 Label

Many biological annotations are organized into hierarchies of abstract concepts, known as ontologies [8]. Each ontology term (a node in the ontology graph) is associated with a set of genes provided by current curated knowledge. By the hierarchical nature of the ontologies, if a gene is associated with an ontology term, then it must also be associated with any more general ontology terms. Thus, we first propagate gene annotations over the ontology, resulting in highly redundant gene set collection. Then, we pick a non-redundant subset of the gene sets with other filtering criteria to set up the final labeled data in the form of **Y** ∈ {0, 1} *N ×p* with *N* number of genes and *p* number of labels (Table 1b).

**Table 1:** Main dataset statistics. The network density is computed as the ratio of existing edges over all possible undirected edges: 2(#*edges*)*/*(#*nodes ×* (#*nodes −* 1)).

| (a) Network statistics |  |  |  | (b) Label set collection statistics |  |  |
| --- | --- | --- | --- | --- | --- | --- |
|  | # nodes | # edges | Density |  | # tasks | # positives |
| BioGRID | 18,951 | 1,103,298 | 0.0030 | DisGeNET | 298 | $198.6 \pm 135.0$ |
| HumanNet | 17,211 | 847,104 | 0.0029 | GOBP | 105 | $88.4 \pm 35.3$ |

#### 2.2.1 Annotated ontology

An ontology is a directed acyclic graph *H* = (*V*_*H*_, *E*_*H*_), where *v* ∈ *V*_*H*_ is an ontology term. The directed edge (*u, v*) ∈ *E*_*H*_ indicates *v* is a parent of *u*, that is, *v* is more abstract or general concept containing *u*. Consider *B* as the set of all genes we are interested in, and *P*(*B*) be the power set of *B*. Given a gene annotation data, represented as a set of pairs 𝒜 = {(*b, v*)_*i*_}, where *b ∈ B* and *v ∈ V*_*H*_ are gene and ontology term, we represent the *raw annotation J*_0_ : *V*_*H*_ → 𝒫(*B*) as a map from an ontology term to a set of genes, so that *J*_0_(*v*) = {*b* : (*b, v*) ∈ 𝒜 }. We then propagate the *raw annotation J*_0_(*·*) over the ontology *H* into a *propagated annotation J*(*·*), where *J*(*v*) = ∪_*v*′:∃ path from *v* ′ to *v*_ *J*_0_(*v*′), as depicted in Figure 1b.

##### Downloading

All the ontology data is downloaded from the OBO Foundry [79] (Figure 1a), which actively maintains tens of large-scale biological ontologies. In the first release, we focus on three different annotations, Gene Ontology (GO) [4], DisGeNET [74], and DISEASES [30]. We naturally split GO into three different sub-collections, namely biological process (GOBP), cellular component (GOCC), and molecular function (GOMF), resulting in a total number of five label collections

#### 2.2.2 Filtering

The propagated annotations contain highly redundant gene sets, which may bias the benchmarking evaluations toward genes sets that commonly appear. To address this, we adapt the non-redundant gene set selection scheme used in Liu et al. [54]. In brief, we construct a graph of gene sets based on the redundancy between two gene sets, and recursively extract the most representative gene set in each connected component (Figure 1b). In addition, we provide several other filtering methods, such as filtering gene sets based on their sizes, number of occurrences, and their existence in the gene interaction network, to help further clean up the gene set collection to be used for final evaluation. Advanced users can alter these filtering functionalities to flexibly set up a custom gene set collection to better suit their biological interests besides using the default filtering steps provided by `obnb`.

### 2.3 Data splitting

A rigorous data splitting should closely mimic real-world scenarios to provide accurate and unbiased estimations of the models’ performance. One stringent solution is temporal holdout, where the training input and label data are obtained before a specific time point, and the testing data is only available afterwards [18, 54]. In practice, this temporal setting is often too restrictive and leads to insufficient tasks for evaluation [54]. Thus, by default, we consider a closely related but less strict strategy called *study-bias holdout*.

#### 6/2/2 study-bias holdout

We use top 60% of the most studied genes with at least one associated label for training. The extend to which a gene is studied is based on its number of associated PubMed publications retrieved from [NCBI](https://www.ncbi.nlm.nih.gov/). The 20% least studied genes are used for testing, and the rest of the 20% genes are used for validation. Notice that by splitting up genes this way, some of the tasks may get very few positive examples in one of the splits. Hence, we remove any task whose minimum number of positive examples across splits falls below a threshold value (set to 10 by default). For completeness, we also provide functionalities in `obnb` to generate random splits.

### 2.4 Dataset construction

The network (Section 2.1) and label (Section 2.2) modules provide flexible solutions for processing and setting up datasets. Meanwhile, we provide a default dataset constructor that utilizes the above modules with reasonable settings to set up a dataset given particular selections of network and label. The default dataset construction workflow is as follows.

1. Select the network and label (task collection) of interest.
2. Remove genes in the task collection that are not present in the gene interaction network.
3. Remove tasks whose number of positive examples falls below 50 after the gene filtering above.
4. Setup 6/2/2 study-bias holdout (Section 2.3).

## Experimental settings

To provide solid baselines for future references, we benchmark a wide range of graph ML methods on our OBNB datasets. Due to space limits, we only present a subset of benchmarking results in the main paper and refer readers to Appendix C for all other results. Specifically, we only include datasets generated using the BioGRID or HumanNet network, combined with the GOBP or DisGeNET labels, resulting in four datasets. The two networks were chosen for their distinct characteristics: BioGRID is an unweighted graph whose edges indicate protein interaction evidence from high throughput experiments [82], whereas HumanNet is a weighted graph whose edges indicate much more diverse types of interactions such as gene coexpression and associations derived via literature text-mining [42]. Meanwhile, the two selected label collections cover two broad classes of gene classification problems, namely, gene function prediction (GOBP) and disease gene association prediction (DisGeNET). We report the performance as the average test precision over prior (APOP) score across five seeds (see Appendix A.4 for the mathematical definition and the motivation of choosing APOP as our main metric). The statistics about networks and task collections are shown in Table 1.

### 3.1 Baselines

We consider three general types of methods: (1) label propagation [101] that directly propagates label information over the graph, (2) graph embedding that first extracts the vectorial representations of each node in the graph, which are then used to fit a simple classifier, such as logistic regression, and (3) GNN that learns the mapping from each node to its label space end-to-end. All implementation details, including technical descriptions of the featurization, the backbone graph neural network architecture, and hyperparameter settings, can be found in the Appendix A.

#### Graph embeddings

We include three distinct featurizations in the main result: using the rows of the adjacency matrix as the node features (Adj) [54], using the *node2vec* embedding (N2V) [31], or using the Laplacian EigenMap embedding (LapEigMap) [9]. Extended results using SVD, LINE [84], and Walklets [72] are available in the Appendix (Table S6). Each of these features is coupled with an *l*_2_ regularized logistic regression model (LogReg) for downstream prediction.

#### GNN

We include multiple variations of two GNN, GCN [48] and GAT [87], in the main results. Extended results for SAGE [32], GIN [99], and GatedGCN [12] can be found in the Appendix (Table S6). One major challenge in applying GNN to OBNB datasets is the lack of canonical node features. This is unlike networks in other domains, such as citation networks, where node features are naturally defined using the paper content, such as bag of words [34]. To tackle this problem, we experiment with a diverse selection of node featurization strategies, including using the one hot encoded log degree of the node, using *node2vec*, and many others. We provide detailed descriptions for the 15 different feature construction strategies in Appendix A.2.1, with extended results for GCN and GAT in Table S5.

#### Bag of Tricks (BoT)

In addition to optimizing the model architectures, many tricks in model training and feature augmentations have also shown to be key factors for practically good performance in existing benchmarks [34]. Here, we further investigate the usefulness of several popular tricks, including reusing training labels as node features (LabelReuse) [94] and performing post-correction to the predictions at test time via correct and smooth (C&S) [39].

## 4 Results and discussions

### 4.1 Traditional ML methods perform comparably to GNNs overall

Table 2 shows the overall performance for the selected model on the four primary benchmarking datasets (full baseline results in Table S6). Surprisingly, even after a rather extensive hyperparameter search and GNN architecture tuning (Appendix A.3), logistic regression using graph-derived features still performs comparably to GNNs. For example, the *node2vec* achieves the best performance for GOBP prediction when using both BioGRID and HumanNet. Similar results are obtained when using the other networks (Table S6). The superior performance of traditional ML methods over GNNs is in stark contrast with results reported in recent benchmarking studies [78, 23, 34], highlighting the unique characteristics of biological networks and the challenges in modeling them.

**Table 2:** Performance of different methods on the four primary OBNB datasets evaluated in APOP ↑ aggregated over five seeds. **Bold** indicates the best-performing method within the method class (traditional ML or GNN). Green indicates best performing-method across both classes. See Table S7.

| Model | Features | C&S? | BioGRID |  | HumanNet |  |
| --- | --- | --- | --- | --- | --- | --- |
|  |  |  | DisGeNET | GOBP | DisGeNET | GOBP |
| LabelProp | — | × | **0.931 ± 0.000** | 1.885 ± 0.000 | **3.059 ± 0.000** | 3.806 ± 0.000 |
| LogReg | Adj | × | 0.743 ± 0.000 | 2.528 ± 0.000 | 3.053 ± 0.000 | 3.964 ± 0.006 |
| LogReg | N2V | × | 0.836 ± 0.029 | **2.571 ± 0.015** | 2.433 ± 0.029 | **4.036 ± 0.019** |
| LogReg | LapEigMap | × | 0.864 ± 0.002 | 2.149 ± 0.000 | 2.301 ± 0.000 | 3.778 ± 0.001 |
| GCN | LogDeg | × | 0.773 ± 0.035 | 2.022 ± 0.100 | 2.452 ± 0.107 | 3.524 ± 0.061 |
| GCN | LogDeg | ✓ | 1.026 ± 0.013 | 2.201 ± 0.035 | 2.777 ± 0.031 | 3.743 ± 0.015 |
| GCN$^\dagger$ | N2V+LogDeg+Label | ✓ | 1.014 ± 0.020 | 2.411 ± 0.044 | 3.053 ± 0.078 | 3.921 ± 0.045 |
| GCN$^\ddagger$ | N2V+LogDeg+Label | ✓ | 1.012 ± 0.040 | 2.572 ± 0.066 | **3.116 ± 0.017** | 3.812 ± 0.071 |
| GAT | LogDeg | × | 0.552 ± 0.111 | 1.592 ± 0.408 | 2.547 ± 0.207 | 3.571 ± 0.159 |
| GAT | LogDeg | ✓ | 1.002 ± 0.018 | 2.227 ± 0.024 | 3.007 ± 0.037 | 3.876 ± 0.053 |
| GAT$^\dagger$ | N2V+LogDeg+Label | ✓ | 1.037 ± 0.036 | **2.624 ± 0.070** | 3.100 ± 0.031 | 3.908 ± 0.086 |
| GAT$^\ddagger$ | N2V+LogDeg+Label | ✓ | **1.063 ± 0.023** | 2.562 ± 0.070 | 3.065 ± 0.021 | **3.963 ± 0.082** |

$^\dagger$ recommended BoT, $^\ddagger$ recommended BoT with fully tuned GNNs (see Appendix A.3.1 for tuning details)

#### Recommendation of BoT for GNNs

While plain GNNs without BoT yield relatively unsatisfactory performances, carefully crafted BoT can bring significant enhancement to GNNs in our benchmark. First, a systematic benchmark on GCN and GAT with 15 different feature construction reveals that using *node2vec* as node features consistently achieve top performance across datasets (Figure S2). Second, reusing training labels as features also elevates overall performance, most notably for GAT (Figure S3). Third, C&S post-processing universally improves performance for both GNN and logistic regression methods, with, on average, 0.27 improvement in test APOP scores (Figure S3,S4). In light of these observations, we recommend an optimized BoT as follows: (1) use a combination of LogDeg encodings, *node2vec* embeddings, and training labels as input node features, (2) apply C&S post-processing to refine predictions further. Our results indicate that the recommended BoT helps improve GNNs’ performances by 0.36 APOP test scores on average across datasets.

#### Different models have their own strength

Despite the overall rankings of different methods indicating the superiority of their gene classification capabilities, no single method can achieve the best results across all tasks. We demonstrate this in Figure 2, which shows that the performance of most methods does not correlate with each other across different tasks on the BioGRID-DisGeNET dataset. For example, while LogReg+Adj performs better than GAT+LogDeg overall on the BioGRID-DisGeNET dataset, GAT+LogDeg achieves significantly better predictions (t-test p-value *<* 0.01) for a handful of tasks (Appendix S8), such as iris disorder (MONDO:0002289) and ocular vascular disorder (MONDO:0005552).

**Figure 2.**
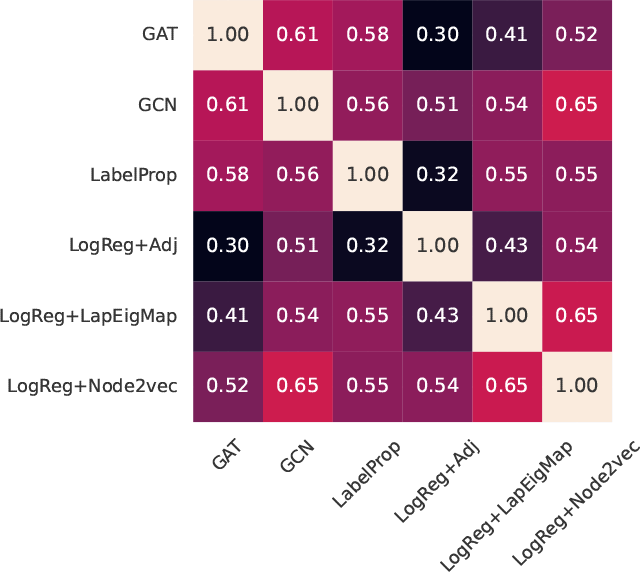
Correlations between methods across tasks in BioGRID-DisGeNET.

Focusing on the optimal model for one or a few tasks of interest is utterly essential. This is because, in practice, experimental biologists often come with only one or a few gene sets and want to either obtain new related genes or reprioritize them based on their relevance to the whole gene set [61]. Therefore, understanding the characteristics of the task that for which a particular model works well over others is the key to making better architectural decisions for a new task of interest and, ultimately, designing new specialized GNN for network biology.

#### Homophily is a driving factor for good predictions

*Homophily* describes the tendency of a node’s neighborhood to have similar labels to itself. Such effects are prevalent in many real-world graphs, such as citation networks, and have been studied extensively recently in the GNN communities as a way to understand *what* information can or cannot be captured by GNN effectively [58, 60]. Intuitively, homophily aligns well with the *guilt-by-association* principle in network biology, which similarly states that genes that interact with each other are likely functionally related. Despite this clear connection, existing homophily measures, such as the homophily ratio [60], do not straightforwardly apply to the OBNB datasets for two reasons.

1. Most established work on homophily consider *multiclass* classification task, where each node has exactly one class label, contrasting with the *multilabel* classification tasks in OBNB.
2. Label information in OBNB datasets is incredibly sparse. On average, there are only hundreds of positive genes per class out of the total number of 20K genes.

One attempt to resolve the first issue is to compute the average homophily ratio for each class independently. However, due to the label scarcity, the resulting metric can not be readily interpreted. In particular, the highest homophily ratio observed in our main benchmarking study is about 0.2 (Figure 3). Datasets with this value are typically categorized into heterophily (not homophily), contradicting the guilt-by-association principle. To address this inconsistency, we propose a corrected version of the homophily ratio that takes label scarcity into account (see Appendix A.5). The main idea is to measure homophily as a relative measurement: *how much more likely nodes labeled in class i are connected with nodes also labeled in class i than those not in class i*? This measure is directly interpretable as it is the log fold change of the homophily between positive and negative examples.

**Figure 3.**
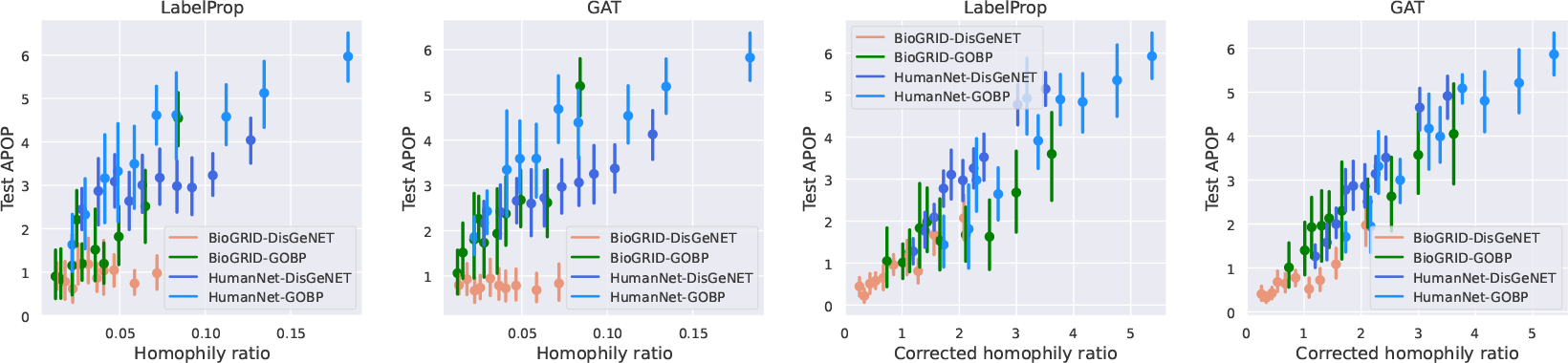
Relationship between the (corrected) homophily ratios of tasks and their performances.

#### Corrected homophily ratio characterizes prediction performance of graph ML methods

As shown in the right two panels of Figure 3, the corrected homophily ratio is well correlated with the prediction performance across different networks and tasks. This positive correlation unveils the fundamental principle that graph ML methods rely on: the underlying graph must provide sufficient contrast between the neighborhoods of the positive and negative examples. Conversely, the left two panels of Figure 3 indicate that the standard homophily ratio fails to adequately account for the performance nuances in low homophily tasks As a result, OBNB opens up new research opportunities in understanding the effects of homophily in GNN by introducing a new type of homophily that differs significantly from those traditionally studied [60].

#### Graph based augmentation tricks offer most advantages on intermediate homophilic tasks

We next investigate the relationship between the corrected homophily ratio and the performance difference between a pair of methods by asking: *is one method systematically better than another method on a particular range of homophily?* The first and second rows of Figure S5 indicate that GNN augmented with different grpah-based tricks, such as C&S and *node2vec* features, provide most apparent advantage at intermediate homophily (corrected homophily ratio of ∼2.5). However, as the homophily increases further, the performance differences diminish, suggesting that all methods would perform equally perfectly as the graph provide more clear clues about the labels. Similar trends are observed for the comparison between GAT+BoT and baseline label propagation and logistic regression methods (Figure S5 third row), where GAT+BoT most significantly outperforms the baselines at intermediate homophily.

### 4.4 Challenges for the OBNB benchmarks and potential future directions

Our systematic benchmarking study paves the way for numerous subsequent explorations. Below, we outline the primary challenges associated with OBNB and suggest potential areas for future research.

#### Challenge 1. Lack of canonical node features

Traditional network biology studies solely rely on biological network structures to gain insights [7, 6, 49]. Meanwhile, the presence of rich node features in many existing graph benchmarks is crucial for the success of GNNs [55], posing a challenge for GNNs in learning without meaningful node features. An exciting and promising future direction for obtaining meaningful node features is by leveraging the sequential or structural information of the gene product (e.g., protein) using large-scale biological pre-trained language models like ESM-2 [53].

#### Challenge 2. Dense and weighted graphs

Many gene interaction networks are dense and weighted by construction, such as coexpression [45] or integrated functional interactions [83]. In more extreme cases, the weighted networks can be fully connected [29]. The networks used in the OBNB benchmark are orders of magnitude denser than citation networks [78] (Table 1a). Thus, scalably and effectively learning from densely weighted gene interaction networks is an area of research to be further explored in the future. One potential solution would be first sparsifying the original dense network before proceeding to train GNN models on them, either by straightforwardly applying an edge cutoff threshold or by using more intricated methods such as spectral sparsification [81].

#### Challenge 3. Scarce labeled data

Curated biological annotations are scarce, posing the challenge of effectively training powerful and expressive models with limited labeled examples. Furthermore, a particular gene can be labeled with more than one biological annotation (multi-label setting), which is much more complex than the multi-class settings in popular benchmarking graph datasets such as Cora and CiteSeer [78]. The data scarcity issue naturally invites the usage of self-supervised [98] graph learning techniques, such as DGI [88], and knowledge transfer via pre-training, such as TxGNN [38]. However, these methods still face the challenge of missing node features (challenge 1).

#### Challenge 4. Low corrected homophily ratio tasks

Our results in Figure 3 show that tasks with low corrected homophily ratios tend to be more challenging to predict, indicating that current tested methods are limited to local information of the underlying graph. This naturally opens up opportunities for designing models that (1) better capture long-range dependencies [22] and (2) exploit higher-order structural information [64].

## Conclusion

We have developed `obnb`, an open-source Python package for rapidly setting up ML-ready bench-marking graph datasets using diverse biomedical networks and gene annotations from publicly available sources. `obnb` takes care of tedious data (pre-)processing steps so that network biologists can easily set up particular datasets with their desired settings, and graph ML researchers can directly use the ML-ready datasets for model development. We have established a comprehensive set of baseline performances on OBNB using a wide range of graph ML methods for future reference and pointed out potential improvements that could further enhance performances. Together, OBNB will help accelerate the development of advanced graph ML methods in network biology toward furthering our understanding of the complex genetic basis of human traits and diseases.

## A Additional information

### A.1 Implementation information

All benchmarking experiments are run using computational nodes with five CPUs, 45GB memory, and a Tesla V100 GPU (32GB).

**Figure S1:**
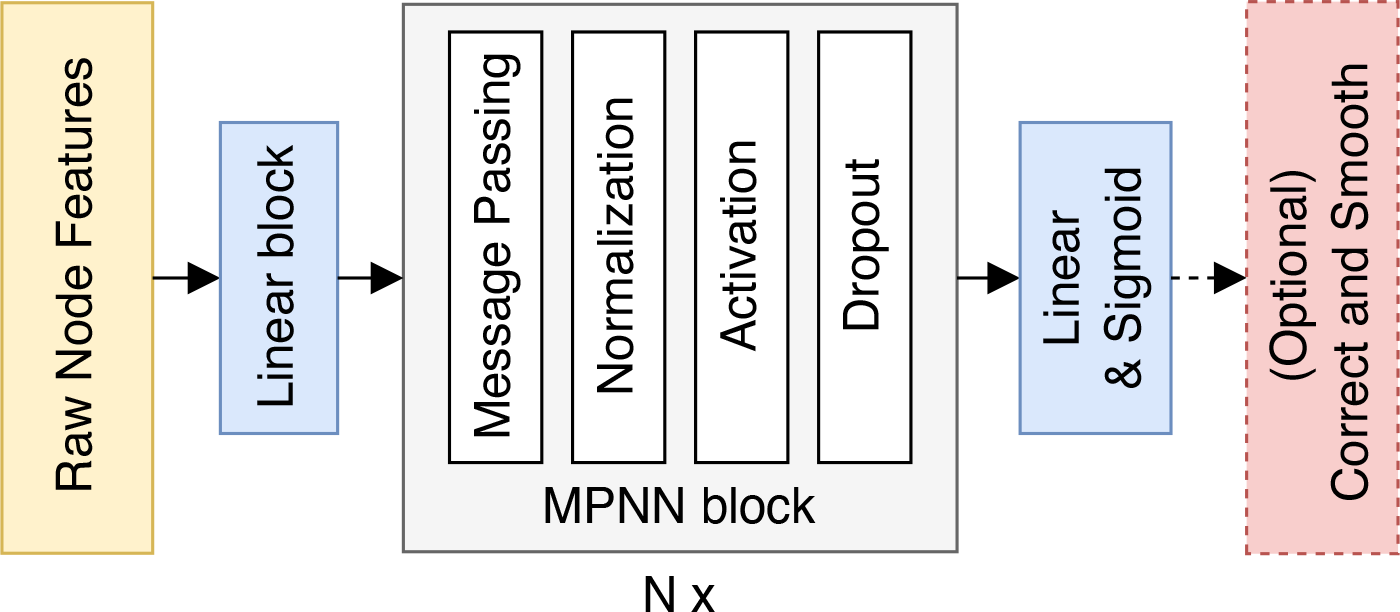
Model architecture overview The basic GNN architecture we used contains the following components:

### A.2 GNN backbone design

#### Feature encoder

Since the genes do not come with canonical features, we need to derive initial node features for the GNN model. The default option we used is OneHotLogDeg with 32 bins (A.2.1). The raw features are first processed by a custom batch norm layer that is activated during testing in addition to training. Then, the processed features are projected into the hidden dimension (*d* = 128 by default) of the model via a linear layer, followed by batch norm and ReLU activation.

#### Message passing layers

We use five message passing layers in the GNN model by default. Each message passing layer contains a convolution block (A.2.2), followed by normalization, non-linear activation, and dropouts. We apply residual connection by summing up the input and the output of the graph convolution block.

#### Prediction head and post-processing

Finally, we apply a linear layer to project the hidden dimension to the dimension matching the number of tasks and apply a sigmoid activation to turn the prediction into binary prediction probabilities. Optionally, we apply correct and smooth (C&S) [39] at test time to correct the predicted probabilities via a two-step label propagation (A.2.3).

##### A.2.1 Node feature design

In this section, we provide an overview of a diverse selection of node feature constructions we used in our benchmark. By default, any initial node feature will be 128-dimensional unless otherwise specified. We start by designing a collection of simple statistics that can be easily derived.

**Constant** uses a one-dimensional trivial feature for every node in the graph.

**RandomNormal** samples 32-dimensional features for each node independently via a standard normal distribution.

**OneHotLogDeg (short for LogDeg)** first computes the log degree of each node in the graph and then uniformly bins the nodes into one of 32 bins based on their log degree. The one-hot encoded node degrees approach has recently been shown to be a great structure encoder, whose utilization can sometimes result in performance superior to using the original node features associated with the graph [17, 55]. Meanwhile, the design choice of using log-uniform grids stems from the scale-free nature of biological networks [2].

**RandomWalkDiag** is the landing probability of a node back to itself after *k* hops, which is also commonly referred to as the random walk structural encodings (RWSE) [21]. It has been used widely in many graph transformer models due to its expressiveness in capturing graph structure [52]. Next, we consider several node feature options derived directly from the adjacency matrix.

**Adj** uses the rows of the adjacency matrix as the node feature. It has been shown previously [54] that logistic regression using the adjacency matrix produced better prediction performance than the commonly used label propagation algorithm for diverse gene classification problems.

**RandProjGaussian and RandProjSparse** are random projections of the adjacency matrix using two different but related approaches. We use the scikit-learn [68] implementations (`GaussianRandomProjection` and `SparseRandomProjection`) to compute these features.

**SVD** uses the left singular vectors of the adjacency matrix as the node features.

**LapEigMap [9]** uses the (*ℓ*_2_ normalized) eigenvectors of the symmetric normalized graph Laplacian as the node features.

Node embeddings [66] are powerful approaches to extracting vectorial representations of each node in a graph and have shown promising results in many biomedical application [54, 100, 5]. Thus, we include a few popular and good-performing node embedding options in our benchmark.

**LINE1 and LINE2 [84]** extracts first- and second-order proximity information from the graph to train the underlying embeddings. We use the GraPE [15] implementation to compute the LINE embeddings.

**Node2vec [31]** extracts node representation using word2vec [63] on node sequences sampled from the graph via biased random walks. The biased random walks, in contrast to an earlier work, DeepWalk [71], which uses an unbiased search, allow the searching strategy to be more flexible, mimicking either breadth-first search or depth-first search in the random walk phase.

**Walklets [72]** is similar to botch Node2vec and DeepWalk in that it runs word2vec using random walks sampled from the graph. However, wallets sample node pairs from the random walk with a specific number of hops, allowing a more explicit control of the multiscale relationships between nodes.

Finally, we experiment with options that let the model learn the node features freely.

**Embedding** lets the model learn the node features freely.

**AdjEmbBag** is similar to Embedding but with an additional aggregation step that sums up the raw embedding of each node from the central node’s neighborhood.

##### A.2.2 Graph neural network summary

In this section, summarize the five tested GNN models under the message-passing framework [26]. Let 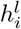 be the *raw* representation of vertex *i* at layer *j*, and 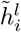 be the corresponding *processed* representation, e.g., after non-linear activation and normalization. The message-passing framework is written as

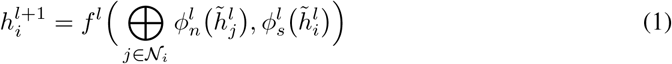

where ⊕ is the aggregation operator, *f* is the update function, *ϕ*_*n*_ and *ϕ*_*s*_ are message functions for neighbors and self. Different GNNs differ in the choices of ⊕, *f, ϕ*_*n*_ and *ϕ*_*s*_ they use.

**GCN [48]** uses a scaled linear transformation as the message function. The scaling factor is computed as the normalized edge weights with added self-loop. The aggregation is done by summing up the transformed message functions.

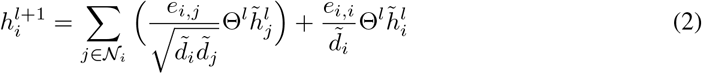

where 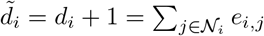 is the degree of vertex *i* with self-loop added.

**SAGE [32]** uses two separate linear transformations as the message functions for neighbors and self. The aggregation is done by taking the sum^1^ of the neighbors’ messages and then adding it to the self-message. We additionally use an affine update function to transform the aggregated messages.

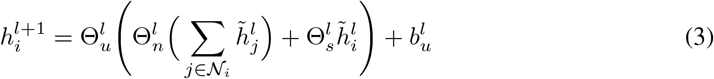

**GIN [99]** uses a multi-layer perceptron (MLP) to update the aggregated messages, which is done by summing up neighbors’ representations and self-representation from the last layer.

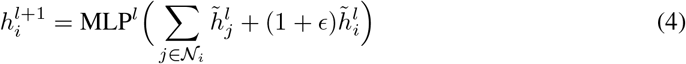

**GAT [87, 13]** uses an attention mechanism to distribute the weights from which the linear transformed messages are aggregated.

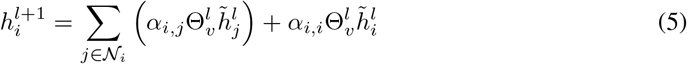

In particular, the original GAT paper [87] formulated the attention scores as follows

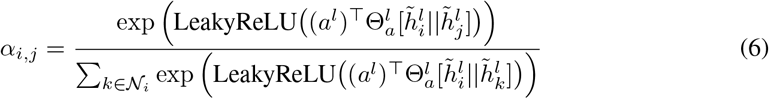

However, it was pointed out in a more recent work, GATv2 [13], that the above formulation was fundamentally limited in the types of attention the model can learn and proposed a correction that showed stronger performances follow.

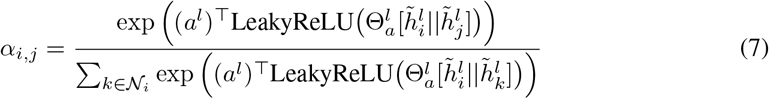

We also observe a slight but significant improvement when using GATv2 attention as opposed to the original GAT. Thus, all reported GAT results are based on v2 attention.

**GatedGCN [12]** uses gating mechanisms to process neighbors’ messages, with linear transformations as the message functions. The aggregation is done by summing up the gated messages from neighbors and adding them to the self-message.

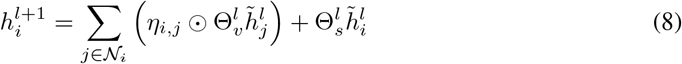

where ⊙ is the elementwise multiplication operator, and the gating coefficients *η*_*i,j*_ are computed as

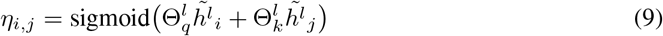

##### A.2.3 Label propagation and C&S

**Label propagation** iteratively propagates the label information from the source nodes outwards through the network neighborhoods. Let **Y** *∈ {*0, 1} ^*n×d*^ be the (training) label matrix, and **M** *∈* R*n×n* be the diffusion operator. The propagated information at step *t*, denoted **F**_*t*_ *∈* R*n×d*, is defined as

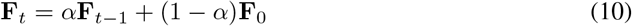

where **F**_0_ = **F** is the features to be propagated, which is the label matrix **Y** in the case of label propagation. The propagated feature is computed by repeating the propagation above until convergence:

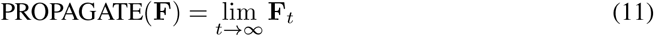

In practice, equation 11 is approximated by applying the propagation until the changes are small enough.

The original label propagation paper [101] uses the symmetric normalized adjacency matrix **D**^*−*^1*/*2 **AD**^*−*^1*/*2 as the diffusion operator, where **A** and **D** are the adjacency matrix and the corresponding diagonal degree matrix. Here, we use the column stochastic matrix **D**^*−*1^**A** as the diffusion operator to resemble the random walk with restart (RWR) implementation that is more commonly used in the network biology literature [16]. We use a propagation parameter *α* of 0.1 (equivalent to a restart parameter of 0.9) for label propagation.

**C&S** implements the idea of using label propagation^2^ to (1) correct for the error made by the model and (2) smooth out the corrected predictions. Specifically, let **E** = **Y**_train_ *−* **Z** be the prediction error matrix, where **Z** is the predicted probabilities resulting from the model. C&S can be summarized as follows.

1. Propagate the error matrix: 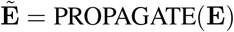.
2. Correct the original prediction with a fixed scale *s:*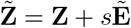.
3. Smooth out the corrected predictions: 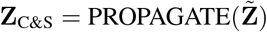

We use *α* = 1.0 for the correction step, *α* = 0.8 for the smoothing step, and a scaling factor of *s* = 1.5.

### A.3 Training and hyperparameter details

All hyperparameters are listed in the configuration files for each model in the benchmarking repository^3^. We summarize the primary default hyperparameters for GNN in Table S1.

For logistic regression baselines, we use the SGD optimizer with a constant learning rate of 200, weight decay of 1e-5, and momentum of 0.9.

#### A.3.1 Fully tuned GNN models

In addition to the baseline GNN experiments where we used the default settings elaborated above, we also tuned GCN and GAT specifically for the four primary benchmarks presented in the main paper with a bag of tricks (BoT). Specifically, we construct node features by concatenating Node2vec embeddings (*d* = 128), OneHotLogDeg encodings (*d* = 32), and LabelReuse. Finally, we apply correct and smooth (C&S) to the GNN predictions as a post-processing step.

We use the Bayesian search hyperparameter optimization strategy provided by Weights&Biases [10] and optimize for the validation APOP scores to tune the dataset-specific hyperparameters for GCN and GAT on primary results presented in Table 2. The search space is listed in Table S2 and the final hyperparameter settings are summarized in Table S3,S4.

### A.4 The log2 fold change of average precision over the prior metric (APOP)

APOP is computed by taking *log*_2_ of the ratio between the average precision and the prior. The prior is computed as the ratio between the number of positives and the total number of samples, which corresponds to the probability that a randomly drawn sample is positive. Thus, APOP indicates how much better (*>* 0), or worse (*<* 0), a prediction is than a random guess, in two-fold change. More precisely,

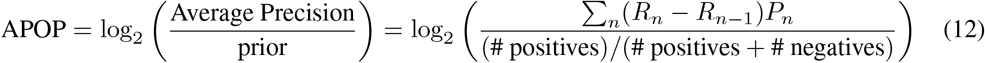

**Table S1:** Default hyperparameter settings.

|  | Parameter | Value |
| --- | --- | --- |
| General architecture | Hidden dimensions | 128 |
|  | Number of layers | 5 |
|  | Activation | ReLU |
|  | Dropout | 0.2 |
|  | Normalization | DiffGroupNorm [103] |
| GAT specific | Heads | 1 |
|  | Attention dropout | 0.05 |
| GIN specific | MLP layers | 2 |
|  | MLP dimensions | 256 |
| | $\epsilon$ | 0.0 |
| Optimizer<br>(AdamW [57]) | Learning rate | 0.01 |
|  | Weight decay | 1e-6 |
|  | Max epochs | 50,000 |
|  | Early stopping patience | 500 |
| Learning rate scheduler<br>(ReduceLROnPlateau) | Scheduler patience | 100 |
|  | Scheduler reduce factor | 0.5 |
|  | Minimum learning rate | 1e-5 |

**Table S2:** Hyperparameter search space for GCN and GAT on the primary datasets.

| Parameter | Search space |
| --- | --- |
| Number of layers | {1, 2, 3, 4, 5, 6, 7, 8} |
| Hidden dimensions | {64, 128, 192, 256} |
| Number of heads (GAT) | {1, 2, 3, 4} |
| Attention dropout (GAT) | {0.0, 0.05, 0.1} |
| Dropout | {0.1, 0.2, 0.3} |
| Normalization | {none, BatchNorm, LayerNorm, PairNorm, DiffGroupNorm} |
| Activation | {ReLU, PReLU, GELU, SELU, ELU} |

**Table S3:** Tuned GCN model on the four primary benchmarks.

|  | BioGRID |  | HumanNet |  |
| --- | --- | --- | --- | --- |
|  | GOBP | DisGeNET | GOBP | DisGeNET |
| Hidden dimension | 192 | 256 | 192 | 64 |
| Number of layers | 3 | 1 | 4 | 4 |
| Activation | SELU | PReLU | GELU | ReLU |
| Dropout | 0.3 | 0.3 | 0.3 | 0.3 |
| Normalization | DiffGroupNorm | DiffGroupNorm | – | DiffGroupNorm |

**Table S4:** Tuned GAT (v2) model on the four primary benchmarks.

|  | BioGRID |  | HumanNet |  |
| --- | --- | --- | --- | --- |
|  | GOBP | DisGeNET | GOBP | DisGeNET |
| Hidden dimension | 128 | 64 | 128 | 64 |
| Number of layers | 4 | 2 | 4 | 8 |
| Number of heads | 1 | 4 | 1 | 1 |
| Attention dropout | 0.05 | 0.1 | 0.1 | 0.1 |
| Activation | SELU | PReLU | PReLU | GELU |
| Dropout | 0.2 | 0.3 | 0.2 | 0.1 |
| Normalization | BatchNorm | DiffGroupNorm | DiffGroupNorm | DiffGroupNorm |

where *R*_*n*_ and *P*_*n*_ are the recall and precision and precision at the *n*^th^ prediction score threshold.

The average precision is also related to the AUPRC^4^, which has been shown to be more suitable than the AUROC^5^ in the case of class imbalance [77, 80]. In our dataset, class imbalance is prevalent, where each class has only one or two hundred of positive examples but thousands of negative examples (Table 1b,S10). Moreover, AUROC is more tolerant of errors made on top predictions and generally only requires that the global distribution of the predictions made is consistent with the true labels. AUPRC and AP, on the other hand, penalize the top predictions’ errors more.

#### Practical relevance

The OBNB benchmarks cover diverse gene prioritization tasks, such as pinpointing relevant genes for a particular disease. This formulation can also be straightforwardly extend to drug recommendation or drug repurposing tasks, by considering known drug targets (genes) as positive examples. For such biomedicine applications, in practice, only a few top predictions made by a predictive method can be experimentally verified by experimentalists due to the high experimental costs. Thus, APOP’s emphasis on top predictions is well-aligned with this practical constraint and encourages methods to make highly accurate top predictions.

### A.5 Corrected Homophily Ratio

Let *G* = (*V, E, w*) be a weighted graph (unweighted graphs can be treated as weighted graphs with identical edge weights), with node set *V*, edge set *E*, and the edge weight function *w* : *V × V →* ℝ. Let *y*_*i*_ : *V → {*0, 1*}p* be the label function for the *i*^th^ class (or label), for *i* ∈ {1, …, *p* }.

We define the set of *labeled nodes V*_labeled_ as the ones that are part of at least one label, i.e., *V*_labeled_ = { *v* ∈ *V* | max_*i* ∈1,…,*p*_ *y*_*i*_(*v*) = 1}. Furthermore, we define the *positively* and *negatively labeled nodes* for the *i*^th^ class as 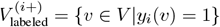 and 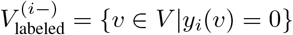, respectively

#### Definition 1 (Node homophily ratio)

*The node homophily ratio of the i*^*th*^ *class for node v is defined as the ratio of its neighborhood that is positively labeled in the i*^*th*^ *class*.

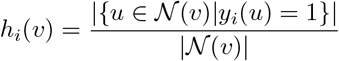

#### Definition 2 (Positive and negative homophily ratio)

*The positive and negative homophily ratios of the i*^*th*^ *class (or label) are defined as the ratio of a node’s neighborhood that are positively labeled for the i*^*th*^ *class averaged over the positively and negatively labeled nodes, respectively*

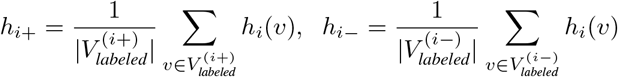

We refer to the *positive homophily ratio* as the *homophily ratio* for short. Note that definition 1 is slightly different from the typical definition of node homophily that is used in previous works targeting multiclass classification settings [69]. Specifically, (1) we only count positive nodes from the neighborhood instead of nodes having the same class as the central node since we are dealing with (multilabel) binary classification, which will also come in handy later for defining the corrected homophily ratio; (2) we only average the node homophily ratios over the positive node sets since our datasets have a notable class imbalance, where most nodes are negatively labeled. This way, the homophily ratio is less skewed towards the majority of negatively labeled nodes.

#### Definition 3 (Corrected homophily ratio)

*The corrected homophily ratio of the i*^*th*^ *class is defined as the expected fold change of node homophily between positively and negatively labeled nodes for class i*

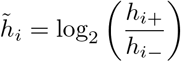

This corrected homophily ratio provides an answer to the following question: *How much more likely are nodes labeled in class i to be connected with other nodes labeled in class i compared to nodes not labeled in class i*? A positive value of the corrected homophily ratio indicates that positive nodes in class *i* have a higher likelihood of interacting with other positive nodes of class *i* than with negative nodes.

### A.6 Community standards and maintenance plans

We follow several community standards to ensure sustainable and maintainable community-wide contributions, which is the key to the continuing improvement of a code base. First, we release the code on GitHub https://github.com/krishnanlab/obnb under the permissive MIT license. Second, we use Sphinx to build the documentation of the code base and host it on ReadTheDocs.org. Several quick-start examples using the package are also provided on the GitHub README page. Third, code quality is ensured via various testing and code-linting automation using tox, pytest, pre-commit hooks, and GitHub actions. Fourth, we provide contribution guidelines and a code of conduct on the GitHub page. Finally, as a part of our commitment to the community, we also put in place a maintenance plan to address GitHub issues, merge pull requests, and release updates periodically to ensure the benchmarks remain adaptive to the evolving needs of the community.

## B Data descriptions

### B.1 Networks

**BioGRID [82] ([MIT License](https://biogrid-downloads.nyc3.digitaloceanspaces.com/LICENSE.txt))** is a protein interaction network curated from primary experimental evidence from the biomedical literature, as well as evidence inferred from low- and high-throughput experiments.

**BioPlex [41] ([Creative Commons Attribution-ShareAlike 4.0 International License](https://bioplex.hms.harvard.edu/explorer/help))** is a protein interaction network whose interactions are measured by affinity purification-mass spectrometry (APMS) analysis shared across two cell lines (HEK293T and HCT116). This shared interaction network encodes core complexes involving many essential proteins that are vital for the cell’s survival.

**ComPPIHumanInt [89] ([CC BY-NC 4.0 License](https://academic.oup.com/nar/article/43/D1/D485/2435307))** is a context-naive version of the cellular compartment-specific ComPPI [89] networks for humans, which are constructed by combining protein interactions from nine different PPI databases (including BioGRID).

**ConsensusPathDB [46] (Free for academic use, see http://cpdb.molgen.mpg.de/ for more Licensing info)** integrates protein interaction evidence (binary protein interaction, protein complex interaction, genetic interaction, metabolic, signaling, etc.) from 31 public databases, in addition to interactions curated from the literature.

**FunCoup [73] ([CC BY-SA 4.0 License](https://funcoup.org/help/))** version 5 is a functional gene interaction network constructed by integrating a wide range of interaction evidence using a redundancy-weighted naive Bayes approach.

**HIPPIE [1] ([CC BY-NC 4.0 License](https://academic.oup.com/nar/article/45/D1/D408/2290937)**) integrates experimentally detected protein interactions from several public databases such as BioGRID.

**HumanBaseTopGlobal [29] ([CC BY 4.0 License](https://humanbase.readthedocs.io/en/latest/))** is the tissue-naive version of the HumanBase tissue-specific gene interaction network collections, which are constructed by integrating hundreds of thousands of publicly available gene expression studies, protein–protein interactions and protein–DNA interactions via a Bayesian approach, calibrated against high-quality manually curated functional gene interactions.

**HuMAP [20] ([CC0 1.0 License](http://humap2.proteincomplexes.org/download))** is a protein interaction network derived from over seven thousand protein complexes by integrating experimental evidence from public resources including AP-MS, large-scale biochemical fraction data, proximity labeling, and RNA hairpin pulldown data.

**HumanNet [42] ([CC BY-SA 4.0 License](https://staging2.inetbio.org/humannetv3/about.php))** is a functional gene interaction network originally designed for disease studies. It contains interaction evidence from gene co-citation from the literature, gene co-expression, pathway, domain profile, genetic interaction, gene neighborhood, phylogenic profile, and other protein interaction data. All these interaction evidence are integrated using a Bayesian statistical framework, resulting in a single value for each pair of genes that indicate the odds ratio for their functional interaction.

**HuRI [59] ([CC BY-4.0 License](http://www.interactome-atlas.org/download))** is a binary protein interaction network constructed via the yeast two-hybrid (Y2H), covering about 90% of the protein-coding genes in the human genome.

**OmniPath [85] (See https://omnipathdb.org/info for Licenses collected for each database)** integrates protein interactions, signaling and regulatory relationships from over 100 resources.

**PCNet [36] ([CC BY-NC 4.0 License](https://www.ndexbio.org/viewer/networks/f93f402c-86d4-11e7-a10d-0ac135e8bacf))** is a protein interaction network constructed by requiring that an edge be supported by at least two out of the 21 selected protein interaction networks, such as BioGRID and ConsensusPathDB.

**ProteomeHD [50] ([CC BY-4.0 License](https://www.ndexbio.org/viewer/networks/4cb4b0f3-83da-11e9-848d-0ac135e8bacf))** is the subnetwork of [50] containing top 0.5% strongest co-regulation signal between pairs of proteins. The co-regulation is measured by proteins’ response to a total of 294 biological perturbations via isotope-labeling mass spectrometry.

**SIGNOR [70] ([CC BY-4.0 License](https://signor.uniroma2.it/about/))** contains manually curated causal signaling relationships between proteins and other biochemical molecules, such as transcriptional activations and phosphorylation.

**STRING [83] ([CC BY-4.0 License](https://string-db.org/cgi/access?footer_active_subpage=licensing))** is a functional interaction network constructed by integrating seven types of gene interaction evidence via a probabilistic approach that calibrates against the KEGG [47] pathway database. The seven evidence types include conserved neighborhood, gene co-occurrence across species, gene fusion events, gene co-expression, other databases, and text-mined interactions.

### B.2 Annotations and Ontologies

**Gene Ontology [4] ([CC BY-4.0 License](https://geneontology.org/docs/go-citation-policy/))** is a structured and standardized system that provides a comprehensive vocabulary to describe the molecular functions (GOMF), biological processes (GOBP), and cellular components (GOCC) associated with genes across different organisms. It aims to unify the representation of biological knowledge and enable effective analysis and interpretation of genomic data.

**Mondo Disease Ontology [86] ([CC BY-4.0 License](https://github.com/monarch-initiative/mondo/blob/master/LICENSE))** is a unifying resource that integrates disease, genotype, and phenotype knowledge across diverse resources, providing a standardized knowledge graph with controlled vocabularies for diseases and phenotypes.

**DisGeNET disease gene annotations [74] ([CC BY-NC 4.0 License](https://academic.oup.com/nar/article/45/D1/D833/2290909))** is a disease gene annotation database that contains a wide range of disease-gene association evidence, including curated annotations, high-throughput experiments and other inferred annotations, animal models, and literature text-mined annotation mostly from BEFREE. By default, we only use the curated and inferred annotations.

**DISEASES disease gene annotations [30]([CC BY License](https://diseases.jensenlab.org/Downloads))** is another disease gene annotation database that has a weekly update schedule for extracting disease gene annotations via text-mining from the fast-growing literature. The text-mined approach uses full text instead of only using the titles and abstracts. Other disease gene annotations from experimental data and other databases are also available.

### B.3 Archived data

In addition to downloading and processing the data directly from the original sources, we also provide archived versions of the data preprocessed by use by running the default OBNB processing pipelines. The archived data is versioned with DOI’s and can be found on Zenodo under the record https://zenodo.org/record/8045270 for extended baseline performance references.

*†* recommended BoT, *‡* recommended BoT with fully tuned GNNs (see Appendix A.3.1 for tuning details)

## C Additional results

**Table S5:** Ablation study on different initial node feature constructions for GCN and GAT. Reported values are average test APOP scores aggregated over five random seeds. The best score within a group (network x label x model) is bolded.

| Network | Model | Feature | DISEASES | DisGeNET | GOBP |
| --- | --- | --- | --- | --- | --- |
| BioGRID | GCN | Constant | 0.373 ± 0.018 | 0.451 ± 0.210 | 0.557 ± 0.144 |
| | | RandomNormal | 1.329 ± 0.043 | 0.867 ± 0.017 | 2.121 ± 0.114 |
| | | OneHotLogDeg | 1.077 ± 0.057 | 0.827 ± 0.050 | 2.011 ± 0.148 |
| | | RandomWalkDiag | 0.844 ± 0.103 | 0.702 ± 0.064 | 1.358 ± 0.044 |
| | | Adj | **1.442 ± 0.120** | 0.899 ± 0.108 | 2.148 ± 0.045 |
| | | RandProjGaussian | 1.288 ± 0.041 | 0.828 ± 0.034 | 2.278 ± 0.084 |
| | | RandProjSparse | 1.264 ± 0.047 | 0.749 ± 0.057 | 2.306 ± 0.088 |
| | | SVD | 1.259 ± 0.076 | 0.808 ± 0.030 | 2.318 ± 0.080 |
| | | LapEigMap | 1.415 ± 0.030 | 0.963 ± 0.045 | 2.378 ± 0.098 |
| | | LINE1 | 1.258 ± 0.041 | 0.798 ± 0.023 | 2.376 ± 0.081 |
| | | LINE2 | 1.312 ± 0.027 | 0.857 ± 0.083 | 2.416 ± 0.048 |
| | | Node2vec | 1.392 ± 0.051 | **0.965 ± 0.055** | **2.487 ± 0.112** |
| | | Walklets | 1.366 ± 0.044 | 0.886 ± 0.079 | 2.438 ± 0.052 |
| | | Embedding | 1.348 ± 0.073 | 0.788 ± 0.060 | 2.147 ± 0.113 |
| | | AdjEmbBag | 1.338 ± 0.071 | 0.798 ± 0.080 | 2.031 ± 0.089 |
| | GAT | Constant | 0.399 ± 0.039 | 0.362 ± 0.071 | 0.398 ± 0.071 |
| | | RandomNormal | 1.290 ± 0.126 | 0.755 ± 0.140 | 2.268 ± 0.047 |
| | | OneHotLogDeg | 0.946 ± 0.289 | 0.803 ± 0.066 | 2.181 ± 0.080 |
| | | RandomWalkDiag | 0.873 ± 0.115 | 0.554 ± 0.160 | 1.702 ± 0.068 |
| | | Adj | 1.290 ± 0.320 | 0.594 ± 0.154 | 1.972 ± 0.230 |
| | | RandProjGaussian | 1.357 ± 0.073 | 0.897 ± 0.070 | 2.402 ± 0.081 |
| | | RandProjSparse | 1.351 ± 0.091 | 0.842 ± 0.063 | 2.461 ± 0.062 |
| | | SVD | 1.019 ± 0.199 | 0.700 ± 0.081 | 2.384 ± 0.058 |
| | | LapEigMap | 1.423 ± 0.072 | **0.914 ± 0.103** | 2.541 ± 0.042 |
| | | LINE1 | **1.480 ± 0.073** | 0.874 ± 0.071 | 2.486 ± 0.040 |
| | | LINE2 | 1.454 ± 0.043 | 0.888 ± 0.057 | 2.463 ± 0.138 |
| | | Node2vec | 1.416 ± 0.126 | 0.877 ± 0.032 | **2.585 ± 0.052** |
| | | Walklets | 1.464 ± 0.072 | 0.852 ± 0.073 | 2.392 ± 0.170 |
| | | Embedding | 1.375 ± 0.092 | 0.834 ± 0.056 | 2.279 ± 0.197 |
| | | AdjEmbBag | 0.645 ± 0.049 | 0.525 ± 0.041 | 1.242 ± 0.062 |
| HumanNet | GCN | Constant | 0.385 ± 0.018 | 0.898 ± 0.873 | 0.610 ± 0.037 |
| | | RandomNormal | 3.006 ± 0.112 | 2.511 ± 0.031 | 3.302 ± 0.104 |
| | | OneHotLogDeg | 2.873 ± 0.134 | 2.552 ± 0.059 | 3.574 ± 0.116 |
| | | RandomWalkDiag | 2.056 ± 0.102 | 1.672 ± 0.171 | 3.001 ± 0.149 |
| | | Adj | **3.562 ± 0.049** | 2.788 ± 0.096 | 3.715 ± 0.077 |
| | | RandProjGaussian | 3.346 ± 0.096 | 2.768 ± 0.036 | 3.654 ± 0.054 |
| | | RandProjSparse | 3.341 ± 0.085 | 2.784 ± 0.021 | 3.759 ± 0.039 |
| | | SVD | 3.211 ± 0.062 | 2.723 ± 0.064 | 3.780 ± 0.075 |
| | | LapEigMap | 3.219 ± 0.077 | 2.676 ± 0.072 | 3.735 ± 0.068 |
| | | LINE1 | 2.883 ± 0.097 | 2.560 ± 0.049 | 3.700 ± 0.078 |
| | | LINE2 | 2.918 ± 0.070 | 2.563 ± 0.082 | 3.656 ± 0.038 |
| | | Node2vec | 3.359 ± 0.070 | **2.798 ± 0.024** | 3.816 ± 0.039 |
| | | Walklets | 3.464 ± 0.066 | 2.762 ± 0.053 | **3.842 ± 0.052** |
| | | Embedding | 3.389 ± 0.051 | 2.637 ± 0.032 | 3.538 ± 0.040 |
| | | AdjEmbBag | 3.499 ± 0.032 | 2.700 ± 0.066 | 3.482 ± 0.048 |
| | GAT | Constant | 0.280 ± 0.043 | 0.449 ± 0.047 | 0.333 ± 0.035 |
| | | RandomNormal | 3.442 ± 0.331 | 2.758 ± 0.204 | 3.535 ± 0.126 |
| | | OneHotLogDeg | 3.052 ± 0.312 | 2.605 ± 0.074 | 3.661 ± 0.088 |
| | | RandomWalkDiag | 2.623 ± 0.862 | 2.340 ± 0.285 | 3.409 ± 0.059 |
| | | Adj | **3.791 ± 0.071** | 2.943 ± 0.053 | 3.770 ± 0.029 |
| | | RandProjGaussian | 3.390 ± 0.105 | 2.749 ± 0.134 | 3.803 ± 0.103 |
| | | RandProjSparse | 3.477 ± 0.135 | 2.811 ± 0.126 | 3.773 ± 0.085 |
| | | SVD | 3.691 ± 0.080 | 2.653 ± 0.101 | 3.792 ± 0.076 |
| | | LapEigMap | 3.438 ± 0.269 | 2.837 ± 0.180 | 3.907 ± 0.034 |
| | | LINE1 | 3.630 ± 0.058 | 2.794 ± 0.163 | 3.857 ± 0.085 |
|  |  | LINE2 | 3.430 ± 0.133 | 2.867 ± 0.051 | 3.779 ± 0.123 |
|  |  | Node2vec | 3.662 ± 0.036 | **2.975 ± 0.027** | **3.947 ± 0.090** |
|  |  | Walklets | 3.483 ± 0.103 | 2.887 ± 0.092 | 3.885 ± 0.112 |
|  |  | Embedding | 3.539 ± 0.227 | 2.797 ± 0.087 | 3.738 ± 0.090 |
|  |  | AdjEmbBag | 3.592 ± 0.086 | 2.802 ± 0.065 | 3.692 ± 0.075 |
| ComPPIHumanInt | GCN | Constant | 0.263 ± 0.026 | 0.307 ± 0.014 | 0.440 ± 0.039 |
|  |  | RandomNormal | 1.544 ± 0.024 | 1.075 ± 0.058 | 2.411 ± 0.064 |
|  |  | OneHotLogDeg | 1.394 ± 0.077 | 0.964 ± 0.043 | 2.244 ± 0.072 |
|  |  | RandomWalkDiag | 0.946 ± 0.109 | 0.759 ± 0.053 | 1.652 ± 0.081 |
|  |  | Adj | 1.471 ± 0.024 | 1.037 ± 0.074 | 2.320 ± 0.049 |
|  |  | RandProjGaussian | 1.527 ± 0.067 | 1.069 ± 0.071 | 2.649 ± 0.030 |
|  |  | RandProjSparse | 1.558 ± 0.069 | 1.030 ± 0.046 | 2.593 ± 0.058 |
|  |  | SVD | 1.459 ± 0.026 | **1.112 ± 0.056** | 2.622 ± 0.091 |
|  |  | LapEigMap | **1.626 ± 0.037** | 1.110 ± 0.061 | 2.484 ± 0.058 |
|  |  | LINE1 | 1.536 ± 0.103 | 1.057 ± 0.049 | 2.540 ± 0.049 |
|  |  | LINE2 | 1.606 ± 0.062 | 1.060 ± 0.041 | 2.588 ± 0.016 |
|  |  | Node2vec | 1.546 ± 0.080 | 1.072 ± 0.047 | **2.744 ± 0.067** |
|  |  | Walklets | 1.564 ± 0.053 | 1.086 ± 0.054 | 2.593 ± 0.034 |
|  |  | Embedding | 1.356 ± 0.040 | 0.863 ± 0.068 | 2.190 ± 0.065 |
|  |  | AdjEmbBag | 1.390 ± 0.042 | 0.929 ± 0.071 | 2.366 ± 0.080 |
|  | GAT | Constant | 0.323 ± 0.052 | 0.327 ± 0.056 | 0.366 ± 0.047 |
|  |  | RandomNormal | 1.386 ± 0.072 | 1.171 ± 0.107 | 2.584 ± 0.079 |
|  |  | OneHotLogDeg | 1.287 ± 0.148 | 0.756 ± 0.272 | 2.197 ± 0.222 |
|  |  | RandomWalkDiag | 1.197 ± 0.097 | 0.811 ± 0.044 | 1.933 ± 0.211 |
|  |  | Adj | 1.348 ± 0.198 | 0.791 ± 0.095 | 2.087 ± 0.035 |
|  |  | RandProjGaussian | 1.565 ± 0.027 | 1.074 ± 0.065 | 2.706 ± 0.107 |
|  |  | RandProjSparse | 1.562 ± 0.067 | 1.107 ± 0.029 | 2.728 ± 0.068 |
|  |  | SVD | 1.382 ± 0.076 | 1.050 ± 0.054 | 2.730 ± 0.058 |
|  |  | LapEigMap | 1.622 ± 0.044 | 1.192 ± 0.043 | 2.742 ± 0.072 |
|  |  | LINE1 | **1.633 ± 0.055** | 1.134 ± 0.071 | 2.625 ± 0.085 |
|  |  | LINE2 | 1.569 ± 0.034 | 1.112 ± 0.067 | 2.656 ± 0.071 |
|  |  | Node2vec | 1.572 ± 0.064 | 1.169 ± 0.079 | 2.766 ± 0.064 |
|  |  | Walklets | 1.623 ± 0.028 | **1.188 ± 0.070** | **2.789 ± 0.043** |
|  |  | Embedding | 1.544 ± 0.036 | 1.011 ± 0.082 | 2.487 ± 0.072 |
|  |  | AdjEmbBag | 0.887 ± 0.078 | 0.611 ± 0.061 | 1.724 ± 0.066 |
| BioPlex | GCN | Constant | 0.342 ± 0.032 | 0.356 ± 0.052 | 0.488 ± 0.083 |
|  |  | RandomNormal | **1.304 ± 0.016** | 0.868 ± 0.030 | 2.445 ± 0.064 |
|  |  | OneHotLogDeg | 1.225 ± 0.065 | **0.909 ± 0.050** | 2.500 ± 0.044 |
|  |  | RandomWalkDiag | 1.242 ± 0.043 | 0.835 ± 0.038 | 2.461 ± 0.138 |
|  |  | Adj | 1.182 ± 0.067 | 0.817 ± 0.023 | **2.613 ± 0.078** |
|  |  | RandProjGaussian | 1.218 ± 0.030 | 0.855 ± 0.066 | 2.583 ± 0.061 |
|  |  | RandProjSparse | 1.256 ± 0.085 | 0.869 ± 0.017 | 2.563 ± 0.053 |
|  |  | SVD | 1.273 ± 0.055 | 0.707 ± 0.046 | 2.513 ± 0.036 |
|  |  | LapEigMap | 1.206 ± 0.038 | 0.785 ± 0.057 | 2.382 ± 0.026 |
|  |  | LINE1 | 1.234 ± 0.028 | 0.790 ± 0.033 | 2.604 ± 0.073 |
|  |  | LINE2 | 1.242 ± 0.059 | 0.789 ± 0.053 | 2.544 ± 0.060 |
|  |  | Node2vec | 1.206 ± 0.042 | 0.784 ± 0.060 | 2.549 ± 0.074 |
|  |  | Walklets | 1.215 ± 0.060 | 0.817 ± 0.045 | 2.558 ± 0.065 |
|  |  | Embedding | 1.157 ± 0.024 | 0.772 ± 0.066 | 2.582 ± 0.034 |
|  |  | AdjEmbBag | 1.240 ± 0.038 | 0.770 ± 0.021 | 2.582 ± 0.100 |
|  | GAT | Constant | 0.275 ± 0.063 | 0.275 ± 0.146 | 0.569 ± 0.143 |
|  |  | RandomNormal | 1.256 ± 0.070 | 0.884 ± 0.047 | 2.489 ± 0.063 |
|  |  | OneHotLogDeg | 1.089 ± 0.100 | 0.793 ± 0.044 | 2.479 ± 0.098 |
|  |  | RandomWalkDiag | 1.041 ± 0.082 | 0.788 ± 0.090 | 2.430 ± 0.085 |
|  |  | Adj | 1.157 ± 0.029 | 0.811 ± 0.042 | 2.582 ± 0.069 |
|  |  | RandProjGaussian | 1.222 ± 0.064 | **0.924 ± 0.041** | 2.588 ± 0.094 |
|  |  | RandProjSparse | 1.213 ± 0.041 | 0.882 ± 0.063 | **2.632 ± 0.009** |
|  |  | SVD | 1.139 ± 0.089 | 0.721 ± 0.073 | 2.539 ± 0.045 |
|  |  | LapEigMap | 1.204 ± 0.062 | 0.832 ± 0.058 | 2.371 ± 0.064 |
|  |  | LINE1 | 1.201 ± 0.060 | 0.802 ± 0.035 | 2.453 ± 0.074 |
|  |  | LINE2 | **1.242 ± 0.034** | 0.850 ± 0.035 | 2.478 ± 0.057 |
|  |  | Node2vec | 1.229 ± 0.039 | 0.768 ± 0.042 | 2.369 ± 0.061 |
| | | Walklets | $1.196 \pm 0.027$ | $0.855 \pm 0.073$ | $2.531 \pm 0.059$ |
| | | Embedding | $1.159 \pm 0.044$ | $0.746 \pm 0.088$ | $2.493 \pm 0.071$ |
| | | AdjEmbBag | $1.183 \pm 0.043$ | $0.784 \pm 0.050$ | $2.522 \pm 0.056$ |
| HuRI | GCN | Constant | $0.346 \pm 0.015$ | $0.529 \pm 0.033$ | $0.384 \pm 0.055$ |
| | | RandomNormal | $0.504 \pm 0.108$ | $0.676 \pm 0.080$ | $0.956 \pm 0.125$ |
| | | OneHotLogDeg | $0.552 \pm 0.080$ | $0.579 \pm 0.076$ | $1.047 \pm 0.102$ |
| | | RandomWalkDiag | $0.549 \pm 0.107$ | $0.484 \pm 0.045$ | $1.027 \pm 0.129$ |
| | | Adj | $0.587 \pm 0.023$ | $0.631 \pm 0.032$ | $0.942 \pm 0.075$ |
| | | RandProjGaussian | $0.676 \pm 0.070$ | $0.673 \pm 0.046$ | $1.016 \pm 0.186$ |
| | | RandProjSparse | **$0.686 \pm 0.087$** | $0.689 \pm 0.055$ | $1.076 \pm 0.101$ |
| | | SVD | $0.628 \pm 0.056$ | $0.667 \pm 0.010$ | $1.005 \pm 0.049$ |
| | | LapEigMap | $0.581 \pm 0.014$ | $0.526 \pm 0.045$ | $0.997 \pm 0.085$ |
| | | LINE1 | $0.658 \pm 0.069$ | $0.690 \pm 0.054$ | $1.053 \pm 0.089$ |
| | | LINE2 | $0.632 \pm 0.125$ | **$0.741 \pm 0.074$** | $1.106 \pm 0.084$ |
| | | Node2vec | $0.566 \pm 0.061$ | $0.738 \pm 0.053$ | $1.126 \pm 0.114$ |
| | | Walklets | $0.596 \pm 0.078$ | $0.681 \pm 0.032$ | **$1.179 \pm 0.056$** |
| | | Embedding | $0.572 \pm 0.070$ | $0.606 \pm 0.034$ | $0.888 \pm 0.109$ |
| | | AdjEmbBag | $0.652 \pm 0.039$ | $0.660 \pm 0.030$ | $1.059 \pm 0.076$ |
| | GAT | Constant | $0.305 \pm 0.078$ | $0.393 \pm 0.077$ | $0.345 \pm 0.060$ |
| | | RandomNormal | $0.524 \pm 0.159$ | $0.541 \pm 0.131$ | $0.968 \pm 0.114$ |
| | | OneHotLogDeg | $0.465 \pm 0.060$ | $0.510 \pm 0.053$ | $0.872 \pm 0.189$ |
| | | RandomWalkDiag | $0.473 \pm 0.057$ | $0.445 \pm 0.129$ | $0.972 \pm 0.057$ |
| | | Adj | **$0.683 \pm 0.109$** | $0.528 \pm 0.053$ | $0.956 \pm 0.157$ |
| | | RandProjGaussian | $0.592 \pm 0.086$ | $0.530 \pm 0.071$ | $1.028 \pm 0.139$ |
| | | RandProjSparse | $0.603 \pm 0.115$ | $0.456 \pm 0.085$ | $1.017 \pm 0.064$ |
| | | SVD | $0.522 \pm 0.099$ | $0.412 \pm 0.029$ | $1.021 \pm 0.099$ |
| | | LapEigMap | $0.599 \pm 0.082$ | $0.557 \pm 0.048$ | $1.071 \pm 0.080$ |
| | | LINE1 | $0.585 \pm 0.068$ | $0.652 \pm 0.043$ | $1.101 \pm 0.033$ |
| | | LINE2 | $0.634 \pm 0.067$ | **$0.743 \pm 0.053$** | $1.095 \pm 0.054$ |
| | | Node2vec | $0.577 \pm 0.017$ | $0.657 \pm 0.067$ | **$1.116 \pm 0.129$** |
| | | Walklets | $0.630 \pm 0.065$ | $0.602 \pm 0.032$ | $1.085 \pm 0.085$ |
| | | Embedding | $0.647 \pm 0.087$ | $0.601 \pm 0.020$ | $0.941 \pm 0.081$ |
| | | AdjEmbBag | $0.603 \pm 0.056$ | $0.581 \pm 0.040$ | $0.902 \pm 0.059$ |
| OmniPath | GCN | Constant | $0.415 \pm 0.083$ | $0.416 \pm 0.044$ | $0.451 \pm 0.046$ |
| | | RandomNormal | $1.417 \pm 0.042$ | $0.910 \pm 0.126$ | $1.798 \pm 0.073$ |
| | | OneHotLogDeg | $1.195 \pm 0.041$ | $0.930 \pm 0.046$ | $1.753 \pm 0.090$ |
| | | RandomWalkDiag | $0.847 \pm 0.038$ | $0.762 \pm 0.045$ | $1.427 \pm 0.056$ |
| | | Adj | **$1.444 \pm 0.019$** | **$1.118 \pm 0.062$** | $2.050 \pm 0.049$ |
| | | RandProjGaussian | $1.381 \pm 0.089$ | $1.014 \pm 0.053$ | $1.935 \pm 0.075$ |
| | | RandProjSparse | $1.338 \pm 0.073$ | $1.066 \pm 0.060$ | $1.935 \pm 0.020$ |
| | | SVD | $1.281 \pm 0.052$ | $0.849 \pm 0.024$ | $1.884 \pm 0.048$ |
| | | LapEigMap | $1.408 \pm 0.030$ | $1.108 \pm 0.092$ | **$2.120 \pm 0.040$** |
| | | LINE1 | $1.366 \pm 0.075$ | $0.946 \pm 0.049$ | $2.027 \pm 0.042$ |
| | | LINE2 | $1.338 \pm 0.065$ | $0.963 \pm 0.054$ | $1.938 \pm 0.096$ |
| | | Node2vec | $1.384 \pm 0.046$ | $1.024 \pm 0.020$ | $1.982 \pm 0.054$ |
| | | Walklets | $1.433 \pm 0.050$ | $1.036 \pm 0.041$ | $2.101 \pm 0.033$ |
| | | Embedding | $1.329 \pm 0.062$ | $0.831 \pm 0.069$ | $1.504 \pm 0.048$ |
| | | AdjEmbBag | $1.373 \pm 0.042$ | $1.085 \pm 0.024$ | $1.917 \pm 0.054$ |
| | GAT | Constant | $0.353 \pm 0.044$ | $0.343 \pm 0.077$ | $0.381 \pm 0.053$ |
| | | RandomNormal | $1.374 \pm 0.041$ | $1.034 \pm 0.091$ | $1.945 \pm 0.104$ |
| | | OneHotLogDeg | $0.775 \pm 0.142$ | $0.773 \pm 0.135$ | $1.685 \pm 0.110$ |
| | | RandomWalkDiag | $0.754 \pm 0.171$ | $0.657 \pm 0.159$ | $1.580 \pm 0.181$ |
| | | Adj | **$1.257 \pm 0.179$** | **$1.184 \pm 0.068$** | $1.865 \pm 0.095$ |
| | | RandProjGaussian | $1.146 \pm 0.117$ | $1.032 \pm 0.041$ | $2.079 \pm 0.126$ |
| | | RandProjSparse | $1.216 \pm 0.075$ | $0.916 \pm 0.121$ | $2.023 \pm 0.272$ |
| | | SVD | $1.032 \pm 0.125$ | $0.889 \pm 0.061$ | $1.916 \pm 0.159$ |
| | | LapEigMap | $1.344 \pm 0.052$ | $1.103 \pm 0.087$ | $2.105 \pm 0.061$ |
| | | LINE1 | $1.263 \pm 0.082$ | $0.936 \pm 0.035$ | $2.077 \pm 0.108$ |
| | | LINE2 | $1.376 \pm 0.041$ | $1.031 \pm 0.104$ | $2.026 \pm 0.081$ |
| | | Node2vec | $1.317 \pm 0.040$ | $1.014 \pm 0.090$ | $2.096 \pm 0.063$ |
| | | Walklets | $1.248 \pm 0.099$ | $1.022 \pm 0.059$ | **$2.190 \pm 0.052$** |
| | | Embedding | $1.371 \pm 0.032$ | $0.917 \pm 0.103$ | $1.738 \pm 0.023$ |
| | | AdjEmbBag | $0.752 \pm 0.076$ | $0.724 \pm 0.140$ | $1.399 \pm 0.171$ |
| ProteomeHD | GCN | Constant | $0.289 \pm 0.112$ | $0.489 \pm 0.021$ | $0.729 \pm 0.481$ |
| | | RandomNormal | $0.695 \pm 0.128$ | $0.583 \pm 0.060$ | $1.267 \pm 0.074$ |
| | | OneHotLogDeg | $0.698 \pm 0.060$ | $0.692 \pm 0.018$ | $1.413 \pm 0.078$ |
| | | RandomWalkDiag | $0.645 \pm 0.072$ | $0.665 \pm 0.053$ | $1.247 \pm 0.049$ |
| | | Adj | $0.656 \pm 0.060$ | $0.588 \pm 0.067$ | $1.386 \pm 0.054$ |
| | | RandProjGaussian | $0.617 \pm 0.096$ | **$0.714 \pm 0.070$** | $1.412 \pm 0.089$ |
| | | RandProjSparse | $0.685 \pm 0.074$ | $0.669 \pm 0.084$ | $1.461 \pm 0.112$ |
| | | SVD | **$0.809 \pm 0.073$** | $0.575 \pm 0.081$ | $1.474 \pm 0.073$ |
| | | LapEigMap | $0.715 \pm 0.017$ | $0.535 \pm 0.090$ | $1.272 \pm 0.035$ |
| | | LINE1 | $0.588 \pm 0.061$ | $0.594 \pm 0.042$ | $1.316 \pm 0.082$ |
| | | LINE2 | $0.646 \pm 0.071$ | $0.630 \pm 0.034$ | $1.323 \pm 0.073$ |
| | | Node2vec | $0.682 \pm 0.049$ | $0.621 \pm 0.071$ | $1.379 \pm 0.102$ |
| | | Walklets | $0.635 \pm 0.093$ | $0.649 \pm 0.082$ | $1.400 \pm 0.141$ |
| | | Embedding | $0.687 \pm 0.074$ | $0.539 \pm 0.024$ | $1.293 \pm 0.158$ |
| | | AdjEmbBag | $0.600 \pm 0.096$ | $0.621 \pm 0.123$ | **$1.480 \pm 0.083$** |
| | GAT | Constant | $0.375 \pm 0.158$ | $0.473 \pm 0.082$ | $0.675 \pm 0.233$ |
| | | RandomNormal | $0.655 \pm 0.115$ | $0.657 \pm 0.129$ | $1.458 \pm 0.049$ |
| | | OneHotLogDeg | $0.705 \pm 0.115$ | $0.687 \pm 0.063$ | $1.339 \pm 0.199$ |
| | | RandomWalkDiag | $0.700 \pm 0.065$ | $0.691 \pm 0.077$ | $1.196 \pm 0.277$ |
| | | Adj | $0.731 \pm 0.149$ | $0.625 \pm 0.068$ | $1.487 \pm 0.080$ |
| | | RandProjGaussian | $0.808 \pm 0.118$ | $0.701 \pm 0.036$ | $1.491 \pm 0.102$ |
| | | RandProjSparse | $0.725 \pm 0.130$ | $0.710 \pm 0.061$ | $1.502 \pm 0.080$ |
| | | SVD | $0.792 \pm 0.137$ | **$0.828 \pm 0.039$** | **$1.556 \pm 0.057$** |
| | | LapEigMap | $0.678 \pm 0.041$ | $0.534 \pm 0.135$ | $1.394 \pm 0.079$ |
| | | LINE1 | $0.638 \pm 0.159$ | $0.627 \pm 0.094$ | $1.338 \pm 0.120$ |
| | | LINE2 | $0.579 \pm 0.079$ | $0.599 \pm 0.115$ | $1.433 \pm 0.076$ |
| | | Node2vec | **$0.839 \pm 0.100$** | $0.702 \pm 0.075$ | $1.446 \pm 0.089$ |
| | | Walklets | $0.644 \pm 0.106$ | $0.653 \pm 0.098$ | $1.544 \pm 0.064$ |
| | | Embedding | $0.650 \pm 0.141$ | $0.659 \pm 0.100$ | $1.460 \pm 0.041$ |
| | | AdjEmbBag | $0.685 \pm 0.064$ | $0.666 \pm 0.079$ | $1.408 \pm 0.052$ |
| SIGNOR | GCN | Constant | $0.388 \pm 0.066$ | $0.425 \pm 0.157$ | $0.500 \pm 0.148$ |
| | | RandomNormal | $1.478 \pm 0.050$ | $1.349 \pm 0.080$ | $1.571 \pm 0.056$ |
| | | OneHotLogDeg | $1.427 \pm 0.032$ | $1.183 \pm 0.087$ | $1.754 \pm 0.066$ |
| | | RandomWalkDiag | $1.268 \pm 0.011$ | $1.022 \pm 0.043$ | $1.601 \pm 0.127$ |
| | | Adj | $1.461 \pm 0.042$ | $1.306 \pm 0.058$ | $1.575 \pm 0.087$ |
| | | RandProjGaussian | $1.488 \pm 0.046$ | $1.253 \pm 0.083$ | $1.764 \pm 0.062$ |
| | | RandProjSparse | $1.535 \pm 0.067$ | $1.369 \pm 0.107$ | $1.780 \pm 0.071$ |
| | | SVD | $1.396 \pm 0.093$ | $1.279 \pm 0.071$ | $1.788 \pm 0.066$ |
| | | LapEigMap | **$1.597 \pm 0.035$** | $1.345 \pm 0.042$ | $1.746 \pm 0.057$ |
| | | LINE1 | $1.546 \pm 0.068$ | $1.354 \pm 0.049$ | $1.813 \pm 0.101$ |
| | | LINE2 | $1.506 \pm 0.084$ | **$1.390 \pm 0.103$** | $1.708 \pm 0.076$ |
| | | Node2vec | $1.566 \pm 0.071$ | $1.345 \pm 0.068$ | **$1.887 \pm 0.115$** |
| | | Walklets | $1.562 \pm 0.076$ | $1.333 \pm 0.088$ | $1.732 \pm 0.049$ |
| | | Embedding | $1.396 \pm 0.035$ | $1.200 \pm 0.076$ | $1.498 \pm 0.064$ |
| | | AdjEmbBag | $1.496 \pm 0.071$ | $1.229 \pm 0.130$ | $1.665 \pm 0.100$ |
| | GAT | Constant | $0.279 \pm 0.038$ | $0.242 \pm 0.051$ | $0.488 \pm 0.176$ |
| | | RandomNormal | $1.603 \pm 0.049$ | $1.281 \pm 0.060$ | $1.707 \pm 0.050$ |
| | | OneHotLogDeg | $1.180 \pm 0.069$ | $1.009 \pm 0.087$ | $1.576 \pm 0.047$ |
| | | RandomWalkDiag | $1.185 \pm 0.115$ | $0.935 \pm 0.151$ | $1.759 \pm 0.115$ |
| | | Adj | $1.538 \pm 0.104$ | $1.251 \pm 0.023$ | $1.529 \pm 0.053$ |
| | | RandProjGaussian | $1.531 \pm 0.085$ | $1.382 \pm 0.107$ | $1.751 \pm 0.037$ |
| | | RandProjSparse | $1.508 \pm 0.033$ | $1.290 \pm 0.073$ | $1.715 \pm 0.090$ |
| | | SVD | $1.529 \pm 0.048$ | $1.178 \pm 0.102$ | $1.718 \pm 0.017$ |
| | | LapEigMap | $1.578 \pm 0.053$ | $1.326 \pm 0.059$ | $1.749 \pm 0.081$ |
| | | LINE1 | $1.583 \pm 0.036$ | $1.282 \pm 0.052$ | $1.844 \pm 0.071$ |
| | | LINE2 | **$1.630 \pm 0.038$** | $1.325 \pm 0.083$ | **$1.863 \pm 0.048$** |
| | | Node2vec | $1.519 \pm 0.066$ | $1.287 \pm 0.047$ | $1.807 \pm 0.033$ |
| | | Walklets | $1.579 \pm 0.045$ | **$1.411 \pm 0.025$** | $1.762 \pm 0.052$ |
| | | Embedding | $1.474 \pm 0.095$ | $1.254 \pm 0.051$ | $1.695 \pm 0.053$ |
| | | AdjEmbBag | $1.457 \pm 0.198$ | $1.154 \pm 0.096$ | $1.405 \pm 0.156$ |

**Figure S2:**
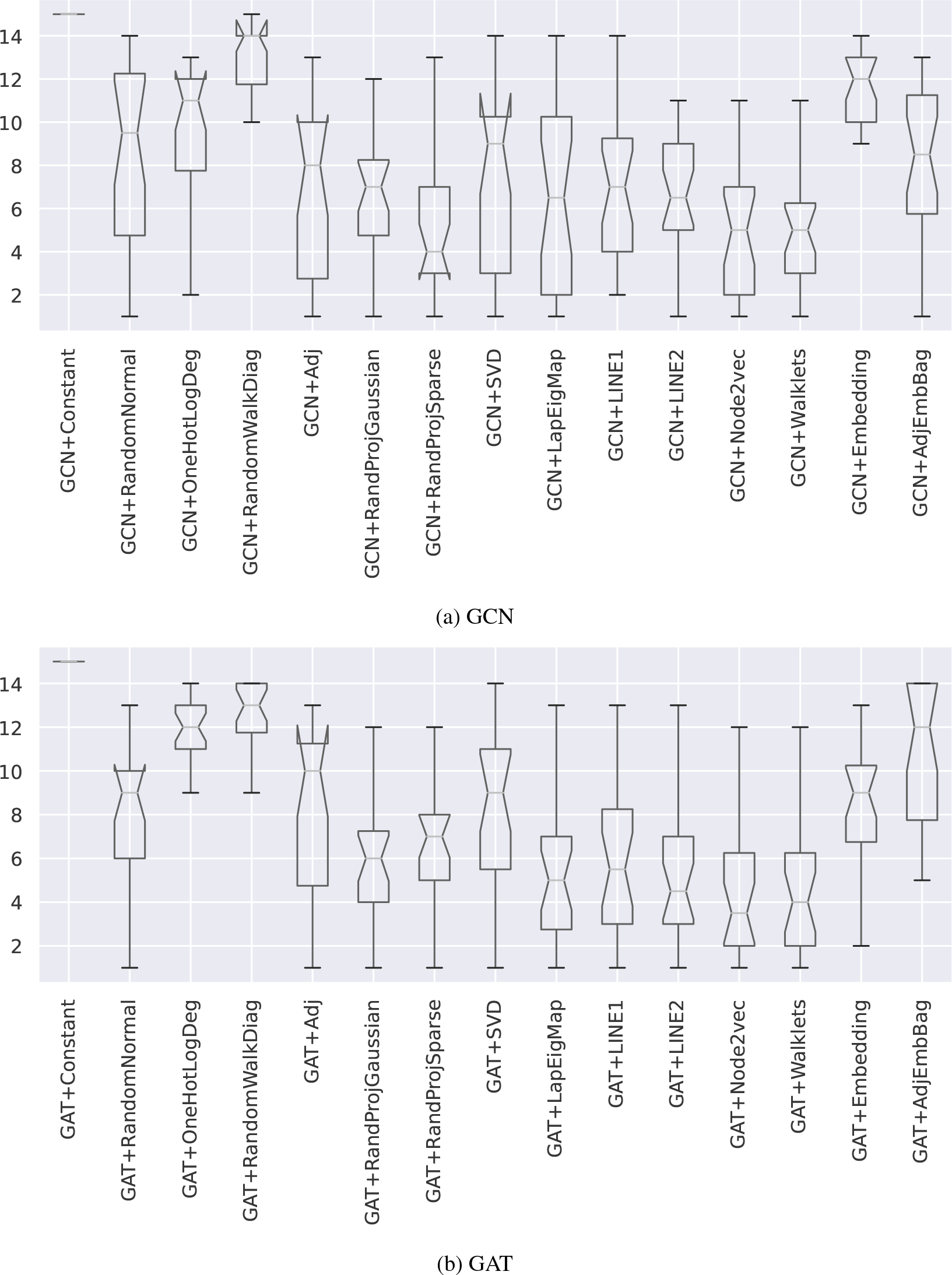
**Boxplots representing the rankings of different feature construction when used by GNNs (lower the better)**. Each point in a box is a ranking of a particular node feature construction on a specific dataset. A lower rank indicates that the particular node feature achieved higher test performance than others. For both GCN and GAT, *node2vec* appears to be the top-ranked node feature overall.

**Table S6:**
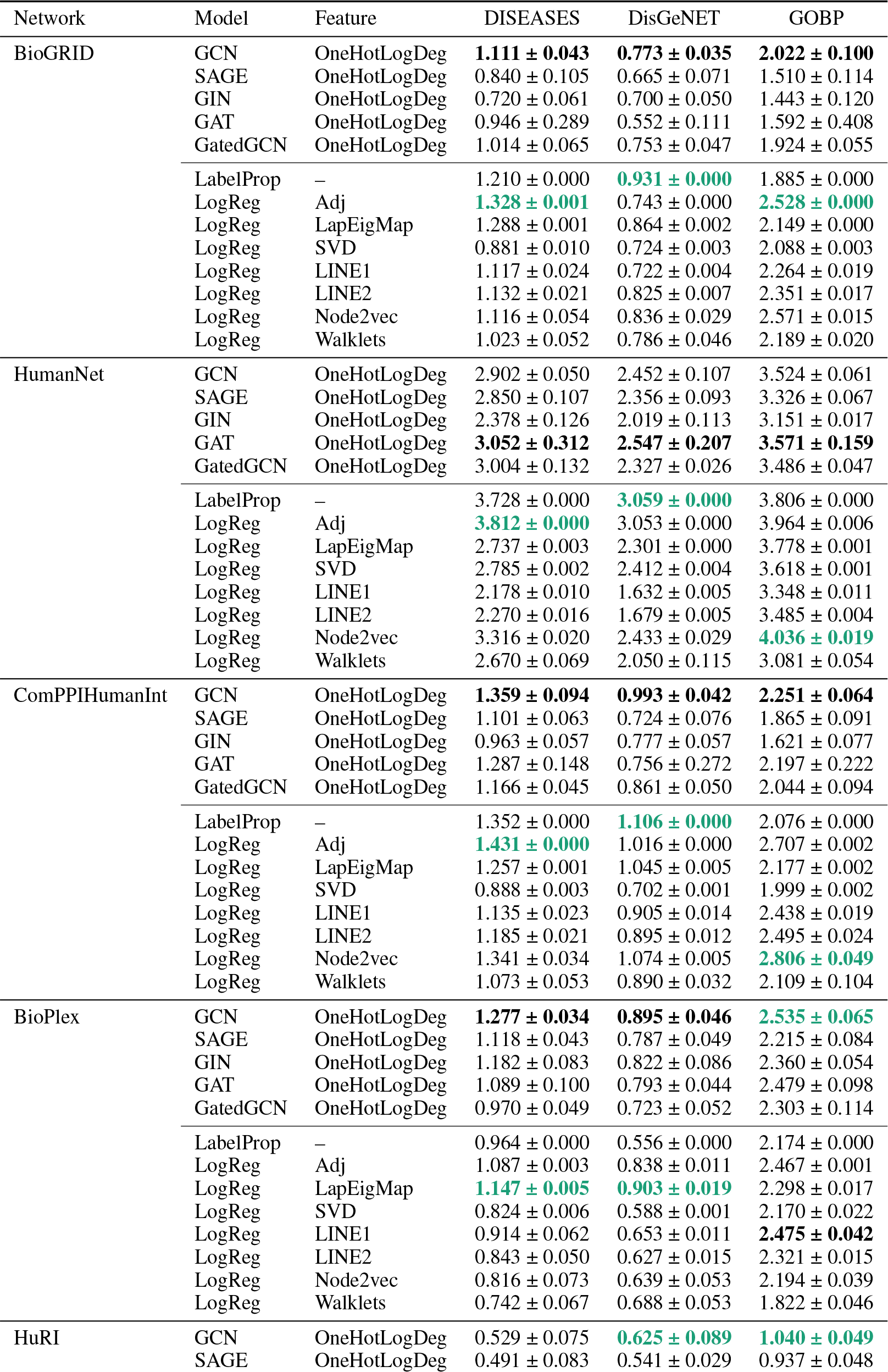
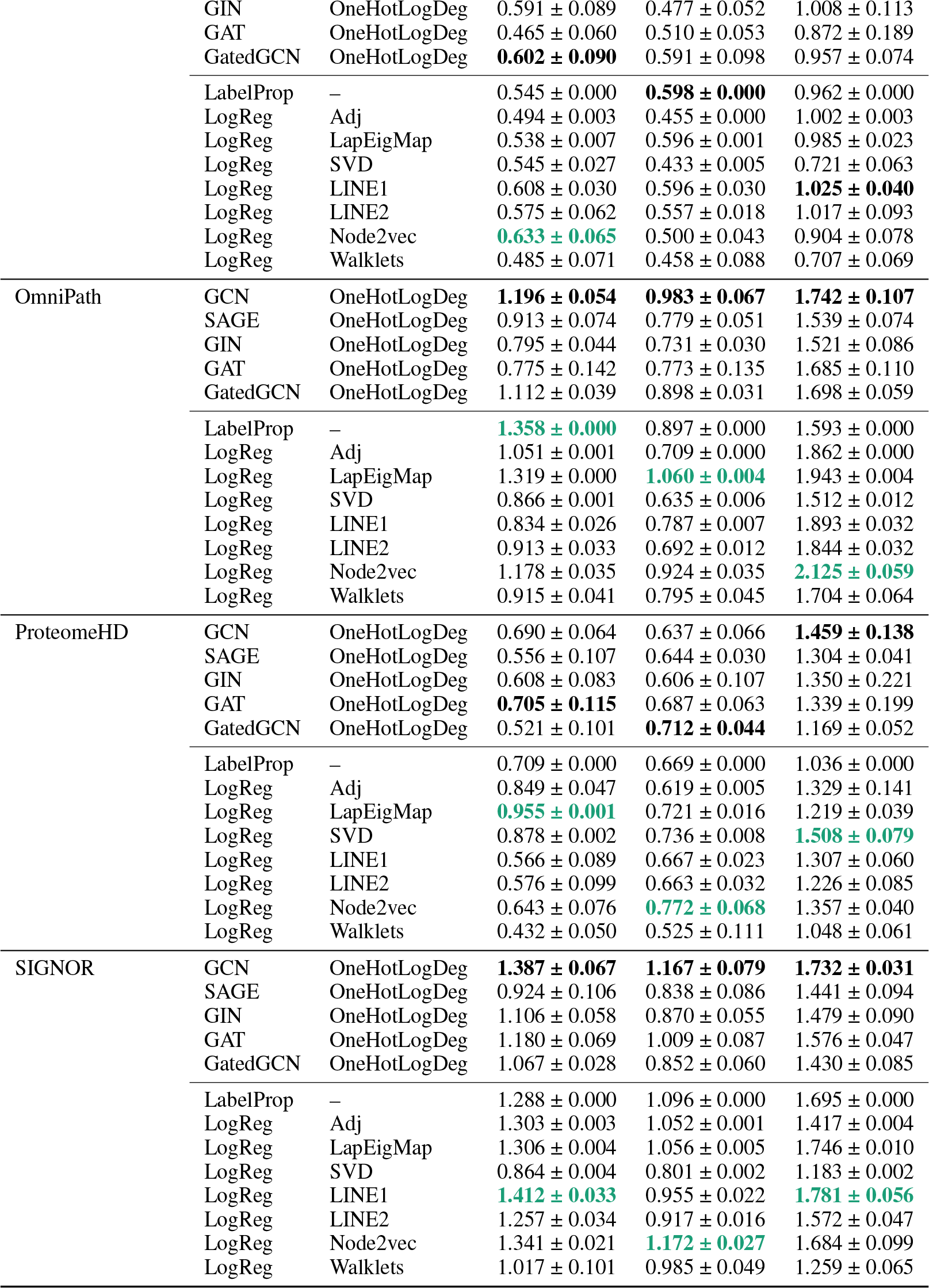
Baseline performance reference. Reported values are average test APOP scores aggregated over five random seeds. The best performance achieved by (1) GNNs or (2) logistic regression and label propagation are **bolded** for each dataset (network *×* label). The best performance across the two methods groups is additionally colored by **green**.

| Network | Model | Feature | DISEASES | DisGeNET | GOBP |
| --- | --- | --- | --- | --- | --- |
| BioGRID | GCN | OneHotLogDeg | **1.111 ± 0.043** | **0.773 ± 0.035** | **2.022 ± 0.100** |
| | SAGE | OneHotLogDeg | 0.840 ± 0.105 | 0.665 ± 0.071 | 1.510 ± 0.114 |
| | GIN | OneHotLogDeg | 0.720 ± 0.061 | 0.700 ± 0.050 | 1.443 ± 0.120 |
| | GAT | OneHotLogDeg | 0.946 ± 0.289 | 0.552 ± 0.111 | 1.592 ± 0.408 |
| | GatedGCN | OneHotLogDeg | 1.014 ± 0.065 | 0.753 ± 0.047 | 1.924 ± 0.055 |
| | LabelProp | – | 1.210 ± 0.000 | **0.931 ± 0.000** | 1.885 ± 0.000 |
| | LogReg | Adj | **1.328 ± 0.001** | 0.743 ± 0.000 | **2.528 ± 0.000** |
| | LogReg | LapEigMap | 1.288 ± 0.001 | 0.864 ± 0.002 | 2.149 ± 0.000 |
| | LogReg | SVD | 0.881 ± 0.010 | 0.724 ± 0.003 | 2.088 ± 0.003 |
| | LogReg | LINE1 | 1.117 ± 0.024 | 0.722 ± 0.004 | 2.264 ± 0.019 |
| | LogReg | LINE2 | 1.132 ± 0.021 | 0.825 ± 0.007 | 2.351 ± 0.017 |
| | LogReg | Node2vec | 1.116 ± 0.054 | 0.836 ± 0.029 | 2.571 ± 0.015 |
| | LogReg | Walklets | 1.023 ± 0.052 | 0.786 ± 0.046 | 2.189 ± 0.020 |
| HumanNet | GCN | OneHotLogDeg | 2.902 ± 0.050 | 2.452 ± 0.107 | 3.524 ± 0.061 |
| | SAGE | OneHotLogDeg | 2.850 ± 0.107 | 2.356 ± 0.093 | 3.326 ± 0.067 |
| | GIN | OneHotLogDeg | 2.378 ± 0.126 | 2.019 ± 0.113 | 3.151 ± 0.017 |
|  | GAT | OneHotLogDeg | **3.052 ± 0.312** | **2.547 ± 0.207** | **3.571 ± 0.159** |
| | GatedGCN | OneHotLogDeg | 3.004 ± 0.132 | 2.327 ± 0.026 | 3.486 ± 0.047 |
| | LabelProp | – | 3.728 ± 0.000 | **3.059 ± 0.000** | 3.806 ± 0.000 |
| | LogReg | Adj | **3.812 ± 0.000** | 3.053 ± 0.000 | 3.964 ± 0.006 |
| | LogReg | LapEigMap | 2.737 ± 0.003 | 2.301 ± 0.000 | 3.778 ± 0.001 |
| | LogReg | SVD | 2.785 ± 0.002 | 2.412 ± 0.004 | 3.618 ± 0.001 |
| | LogReg | LINE1 | 2.178 ± 0.010 | 1.632 ± 0.005 | 3.348 ± 0.011 |
| | LogReg | LINE2 | 2.270 ± 0.016 | 1.679 ± 0.005 | 3.485 ± 0.004 |
| | LogReg | Node2vec | 3.316 ± 0.020 | 2.433 ± 0.029 | **4.036 ± 0.019** |
| | LogReg | Walklets | 2.670 ± 0.069 | 2.050 ± 0.115 | 3.081 ± 0.054 |
| ComPPIHumanInt | GCN | OneHotLogDeg | **1.359 ± 0.094** | **0.993 ± 0.042** | **2.251 ± 0.064** |
| | SAGE | OneHotLogDeg | 1.101 ± 0.063 | 0.724 ± 0.076 | 1.865 ± 0.091 |
| | GIN | OneHotLogDeg | 0.963 ± 0.057 | 0.777 ± 0.057 | 1.621 ± 0.077 |
| | GAT | OneHotLogDeg | 1.287 ± 0.148 | 0.756 ± 0.272 | 2.197 ± 0.222 |
| | GatedGCN | OneHotLogDeg | 1.166 ± 0.045 | 0.861 ± 0.050 | 2.044 ± 0.094 |
| | LabelProp | – | 1.352 ± 0.000 | **1.106 ± 0.000** | 2.076 ± 0.000 |
| | LogReg | Adj | **1.431 ± 0.000** | 1.016 ± 0.000 | 2.707 ± 0.002 |
| | LogReg | LapEigMap | 1.257 ± 0.001 | 1.045 ± 0.005 | 2.177 ± 0.002 |
| | LogReg | SVD | 0.888 ± 0.003 | 0.702 ± 0.001 | 1.999 ± 0.002 |
| | LogReg | LINE1 | 1.135 ± 0.023 | 0.905 ± 0.014 | 2.438 ± 0.019 |
| | LogReg | LINE2 | 1.185 ± 0.021 | 0.895 ± 0.012 | 2.495 ± 0.024 |
| | LogReg | Node2vec | 1.341 ± 0.034 | 1.074 ± 0.005 | **2.806 ± 0.049** |
| | LogReg | Walklets | 1.073 ± 0.053 | 0.890 ± 0.032 | 2.109 ± 0.104 |
| BioPlex | GCN | OneHotLogDeg | **1.277 ± 0.034** | **0.895 ± 0.046** | **2.535 ± 0.065** |
| | SAGE | OneHotLogDeg | 1.118 ± 0.043 | 0.787 ± 0.049 | 2.215 ± 0.084 |
| | GIN | OneHotLogDeg | 1.182 ± 0.083 | 0.822 ± 0.086 | 2.360 ± 0.054 |
| | GAT | OneHotLogDeg | 1.089 ± 0.100 | 0.793 ± 0.044 | 2.479 ± 0.098 |
| | GatedGCN | OneHotLogDeg | 0.970 ± 0.049 | 0.723 ± 0.052 | 2.303 ± 0.114 |
| | LabelProp | – | 0.964 ± 0.000 | 0.556 ± 0.000 | 2.174 ± 0.000 |
| | LogReg | Adj | 1.087 ± 0.003 | 0.838 ± 0.011 | 2.467 ± 0.001 |
| | LogReg | LapEigMap | **1.147 ± 0.005** | **0.903 ± 0.019** | 2.298 ± 0.017 |
| | LogReg | SVD | 0.824 ± 0.006 | 0.588 ± 0.001 | 2.170 ± 0.022 |
| | LogReg | LINE1 | 0.914 ± 0.062 | 0.653 ± 0.011 | **2.475 ± 0.042** |
| | LogReg | LINE2 | 0.843 ± 0.050 | 0.627 ± 0.015 | 2.321 ± 0.015 |
| | LogReg | Node2vec | 0.816 ± 0.073 | 0.639 ± 0.053 | 2.194 ± 0.039 |
| | LogReg | Walklets | 0.742 ± 0.067 | 0.688 ± 0.053 | 1.822 ± 0.046 |
| HuRI | GCN | OneHotLogDeg | 0.529 ± 0.075 | **0.625 ± 0.089** | **1.040 ± 0.049** |
| | SAGE | OneHotLogDeg | 0.491 ± 0.083 | 0.541 ± 0.029 | 0.937 ± 0.048 |
| | GIN | OneHotLogDeg | $0.591 \pm 0.089$ | $0.477 \pm 0.052$ | $1.008 \pm 0.113$ |
| | GAT | OneHotLogDeg | $0.465 \pm 0.060$ | $0.510 \pm 0.053$ | $0.872 \pm 0.189$ |
| | GatedGCN | OneHotLogDeg | **$0.602 \pm 0.090$** | $0.591 \pm 0.098$ | $0.957 \pm 0.074$ |
| | LabelProp | – | $0.545 \pm 0.000$ | **$0.598 \pm 0.000$** | $0.962 \pm 0.000$ |
| | LogReg | Adj | $0.494 \pm 0.003$ | $0.455 \pm 0.000$ | $1.002 \pm 0.003$ |
| | LogReg | LapEigMap | $0.538 \pm 0.007$ | $0.596 \pm 0.001$ | $0.985 \pm 0.023$ |
| | LogReg | SVD | $0.545 \pm 0.027$ | $0.433 \pm 0.005$ | $0.721 \pm 0.063$ |
| | LogReg | LINE1 | $0.608 \pm 0.030$ | $0.596 \pm 0.030$ | **$1.025 \pm 0.040$** |
| | LogReg | LINE2 | $0.575 \pm 0.062$ | $0.557 \pm 0.018$ | $1.017 \pm 0.093$ |
| | LogReg | Node2vec | **$0.633 \pm 0.065$** | $0.500 \pm 0.043$ | $0.904 \pm 0.078$ |
| | LogReg | Walklets | $0.485 \pm 0.071$ | $0.458 \pm 0.088$ | $0.707 \pm 0.069$ |
| OmniPath | GCN | OneHotLogDeg | **$1.196 \pm 0.054$** | **$0.983 \pm 0.067$** | **$1.742 \pm 0.107$** |
| | SAGE | OneHotLogDeg | $0.913 \pm 0.074$ | $0.779 \pm 0.051$ | $1.539 \pm 0.074$ |
| | GIN | OneHotLogDeg | $0.795 \pm 0.044$ | $0.731 \pm 0.030$ | $1.521 \pm 0.086$ |
| | GAT | OneHotLogDeg | $0.775 \pm 0.142$ | $0.773 \pm 0.135$ | $1.685 \pm 0.110$ |
| | GatedGCN | OneHotLogDeg | $1.112 \pm 0.039$ | $0.898 \pm 0.031$ | $1.698 \pm 0.059$ |
| | LabelProp | – | **$1.358 \pm 0.000$** | $0.897 \pm 0.000$ | $1.593 \pm 0.000$ |
| | LogReg | Adj | $1.051 \pm 0.001$ | $0.709 \pm 0.000$ | $1.862 \pm 0.000$ |
| | LogReg | LapEigMap | $1.319 \pm 0.000$ | **$1.060 \pm 0.004$** | $1.943 \pm 0.004$ |
| | LogReg | SVD | $0.866 \pm 0.001$ | $0.635 \pm 0.006$ | $1.512 \pm 0.012$ |
| | LogReg | LINE1 | $0.834 \pm 0.026$ | $0.787 \pm 0.007$ | $1.893 \pm 0.032$ |
| | LogReg | LINE2 | $0.913 \pm 0.033$ | $0.692 \pm 0.012$ | $1.844 \pm 0.032$ |
| | LogReg | Node2vec | $1.178 \pm 0.035$ | $0.924 \pm 0.035$ | **$2.125 \pm 0.059$** |
| | LogReg | Walklets | $0.915 \pm 0.041$ | $0.795 \pm 0.045$ | $1.704 \pm 0.064$ |
| ProteomeHD | GCN | OneHotLogDeg | $0.690 \pm 0.064$ | $0.637 \pm 0.066$ | **$1.459 \pm 0.138$** |
| | SAGE | OneHotLogDeg | $0.556 \pm 0.107$ | $0.644 \pm 0.030$ | $1.304 \pm 0.041$ |
| | GIN | OneHotLogDeg | $0.608 \pm 0.083$ | $0.606 \pm 0.107$ | $1.350 \pm 0.221$ |
| | GAT | OneHotLogDeg | **$0.705 \pm 0.115$** | $0.687 \pm 0.063$ | $1.339 \pm 0.199$ |
| | GatedGCN | OneHotLogDeg | $0.521 \pm 0.101$ | **$0.712 \pm 0.044$** | $1.169 \pm 0.052$ |
| | LabelProp | – | $0.709 \pm 0.000$ | $0.669 \pm 0.000$ | $1.036 \pm 0.000$ |
| | LogReg | Adj | $0.849 \pm 0.047$ | $0.619 \pm 0.005$ | $1.329 \pm 0.141$ |
| | LogReg | LapEigMap | **$0.955 \pm 0.001$** | $0.721 \pm 0.016$ | $1.219 \pm 0.039$ |
| | LogReg | SVD | $0.878 \pm 0.002$ | $0.736 \pm 0.008$ | **$1.508 \pm 0.079$** |
| | LogReg | LINE1 | $0.566 \pm 0.089$ | $0.667 \pm 0.023$ | $1.307 \pm 0.060$ |
| | LogReg | LINE2 | $0.576 \pm 0.099$ | $0.663 \pm 0.032$ | $1.226 \pm 0.085$ |
| | LogReg | Node2vec | $0.643 \pm 0.076$ | **$0.772 \pm 0.068$** | $1.357 \pm 0.040$ |
| | LogReg | Walklets | $0.432 \pm 0.050$ | $0.525 \pm 0.111$ | $1.048 \pm 0.061$ |
| SIGNOR | GCN | OneHotLogDeg | **$1.387 \pm 0.067$** | **$1.167 \pm 0.079$** | **$1.732 \pm 0.031$** |
| | SAGE | OneHotLogDeg | $0.924 \pm 0.106$ | $0.838 \pm 0.086$ | $1.441 \pm 0.094$ |
| | GIN | OneHotLogDeg | $1.106 \pm 0.058$ | $0.870 \pm 0.055$ | $1.479 \pm 0.090$ |
| | GAT | OneHotLogDeg | $1.180 \pm 0.069$ | $1.009 \pm 0.087$ | $1.576 \pm 0.047$ |
| | GatedGCN | OneHotLogDeg | $1.067 \pm 0.028$ | $0.852 \pm 0.060$ | $1.430 \pm 0.085$ |
| | LabelProp | – | $1.288 \pm 0.000$ | $1.096 \pm 0.000$ | $1.695 \pm 0.000$ |
| | LogReg | Adj | $1.303 \pm 0.003$ | $1.052 \pm 0.001$ | $1.417 \pm 0.004$ |
| | LogReg | LapEigMap | $1.306 \pm 0.004$ | $1.056 \pm 0.005$ | $1.746 \pm 0.010$ |
| | LogReg | SVD | $0.864 \pm 0.004$ | $0.801 \pm 0.002$ | $1.183 \pm 0.002$ |
| | LogReg | LINE1 | **$1.412 \pm 0.033$** | $0.955 \pm 0.022$ | **$1.781 \pm 0.056$** |
| | LogReg | LINE2 | $1.257 \pm 0.034$ | $0.917 \pm 0.016$ | $1.572 \pm 0.047$ |
| | LogReg | Node2vec | $1.341 \pm 0.021$ | **$1.172 \pm 0.027$** | $1.684 \pm 0.099$ |
| | LogReg | Walklets | $1.017 \pm 0.101$ | $0.985 \pm 0.049$ | $1.259 \pm 0.065$ |

**Figure S3:**
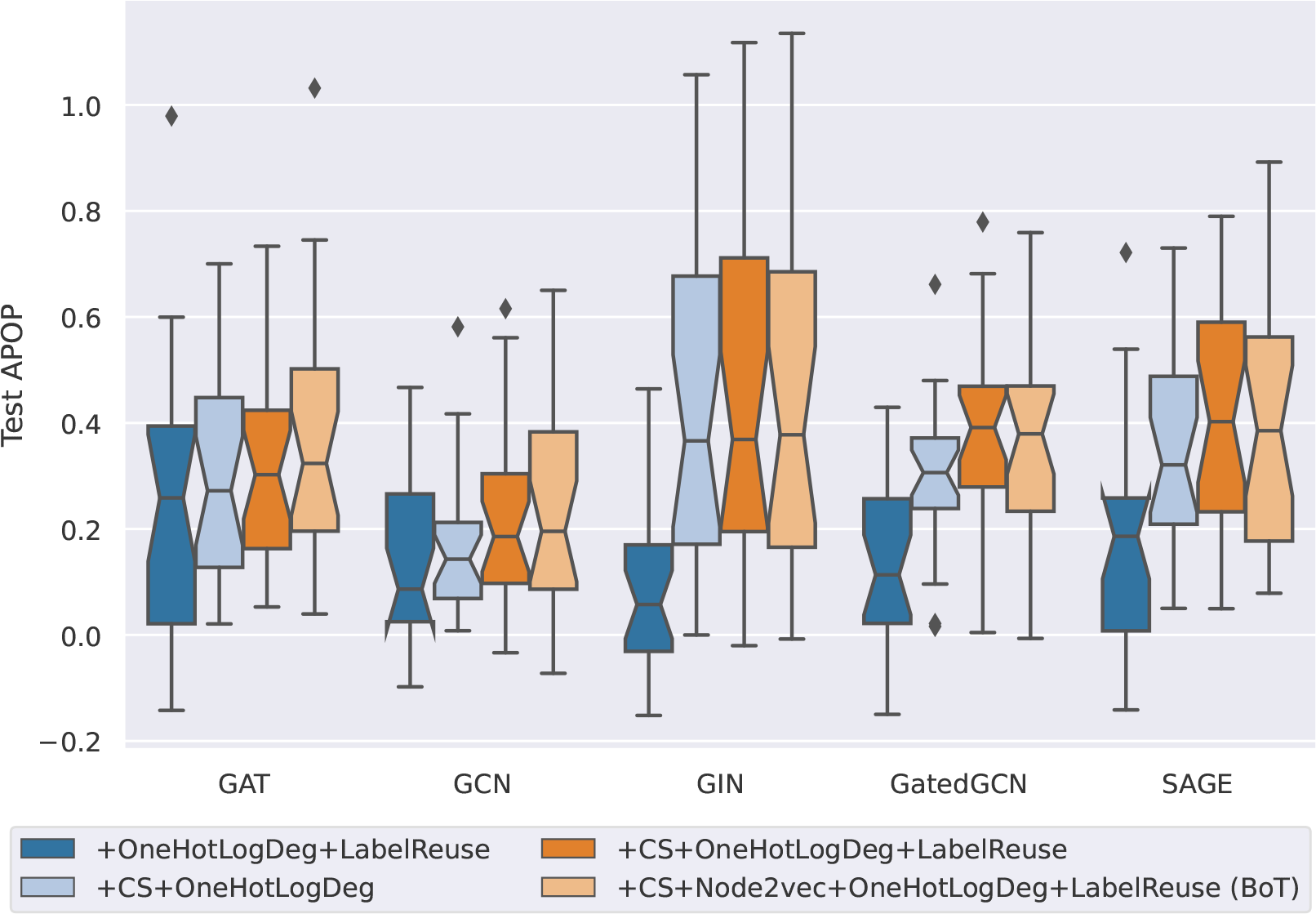
**Boxplots representing the performance improvement of using different tricks with GNNs across datasets**. Each point in a box is the test APOP difference of GNN with added tricks vs. plain GNN using OneHotLogDeg feature on a specific dataset. A positive value implies the added trick improves GNN performance.

**Figure S4:**
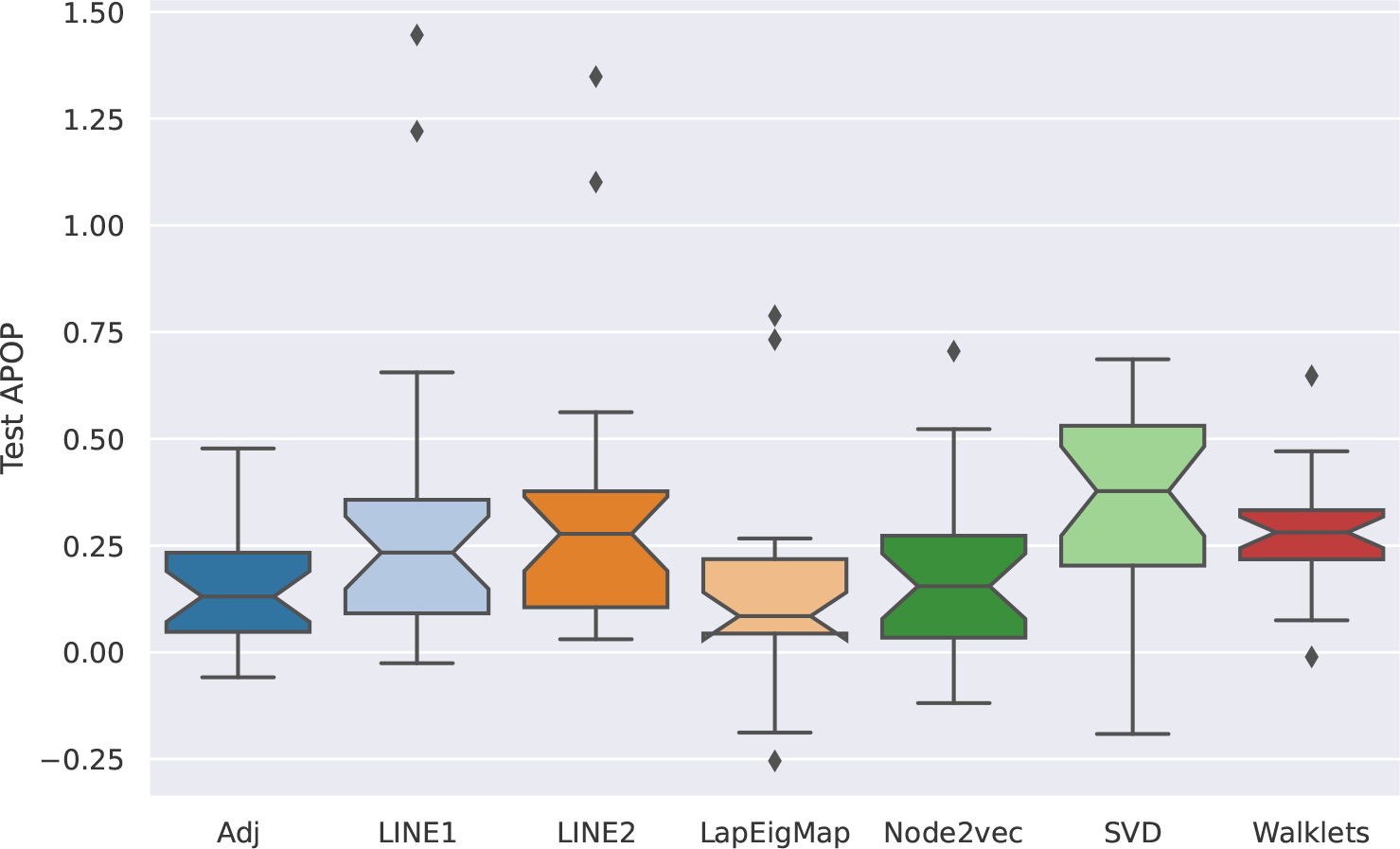
**Boxplots representing the performance improvement of using C&S with logistic regression models across datasets**. Each point in a box is the test APOP difference of logistic regression augmented with C&S vs. plain logistic regression on a specific dataset. A positive value implies C&S improves logistic regression performance.

**Figure S5:**
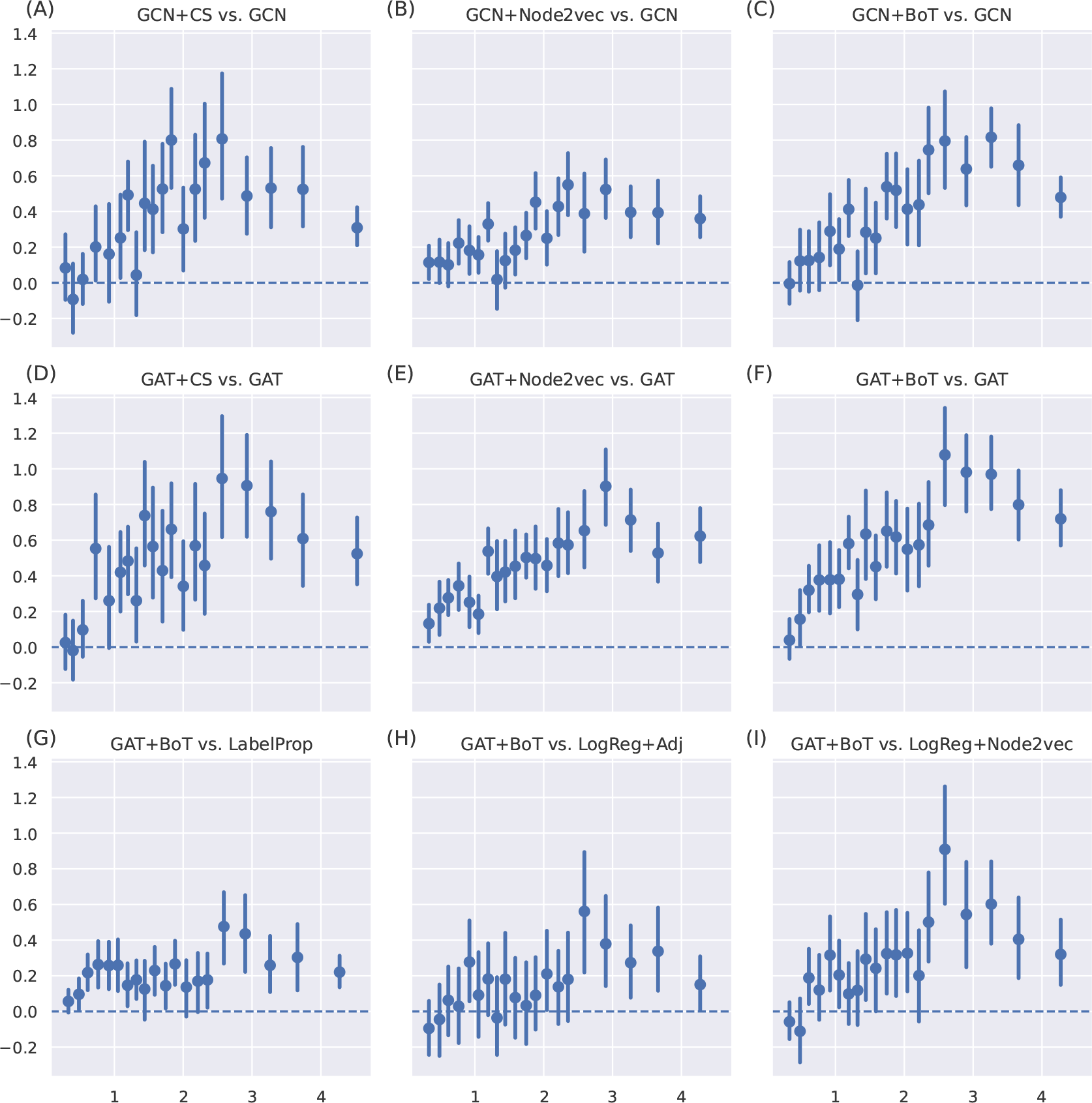
**Relationships between the corrected homophily ratio and performance difference between methods**. Each panel summarizes tasks over three networks (BioGRID, HumanNet, ComPPIHumanInt) and three gene set collections (DISEASES, DisGeNET, GOBP), with a total of 1,672 tasks. In each panel, the x-axis represents the corrected homophily ratio and the y-axis represents the test APOP performance difference between the two methods. The first (A, B, C) and second rows (D, E, F) show the performance improvement to GCN and GAT when augmented with individual tricks or combined BoT. The third row (G, H, I) shows the performance difference between the best GNN method and the baseline methods, including label propagation and logistic regression.

**Table S7:**
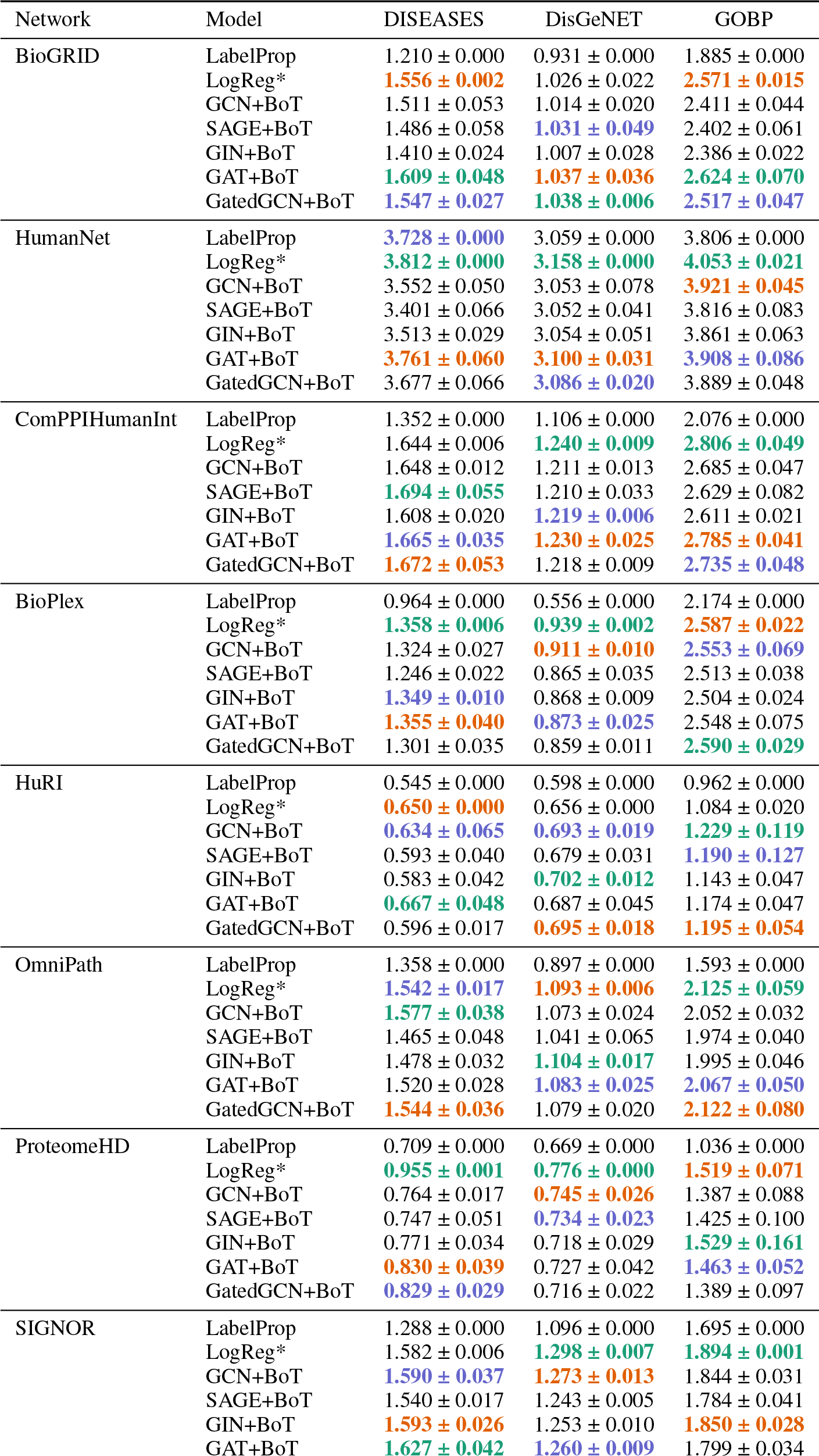
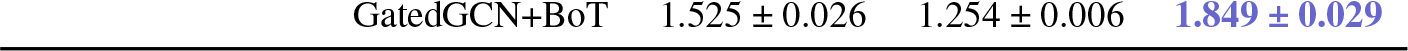
Combined SOTA performance reference. LogReg* are the best test performances achieved from any logistic regression models we have tested for each dataset. Colored texts indicate the **first, second**, and **third** best performance achieved for a particular dataset (network *×* label).

**Table S8:** Task-specific model prediction performance difference on the BioGRID-DisGeNET between GAT and LogReg+Adj.

| Task | GAT | LogReg+Adj | Difference |
| --- | --- | --- | --- |
| MONDO:0002289 | 2.160 | 0.149 | 2.011 |
| MONDO:0001703 | 2.177 | 0.230 | 1.946 |
| MONDO:0001926 | 1.543 | -0.336 | 1.879 |
| MONDO:0005552 | 1.507 | -0.175 | 1.682 |
| MONDO:0018470 | 1.273 | -0.347 | 1.620 |
| ... |  |  |  |
| MONDO:0020018 | 0.699 | 4.487 | -3.788 |
| MONDO:0019743 | 0.773 | 4.876 | -4.103 |
| MONDO:0024757 | 0.316 | 4.443 | -4.127 |
| MONDO:0045011 | 0.264 | 4.619 | -4.355 |
| MONDO:0015160 | 0.280 | 4.828 | -4.549 |

**Table S9:** Statistics for all networks in OBNB (obnbdata-0.1.0)

| Network | Weighted | Num. nodes | Num. edges | Density | Category |
| --- | --- | --- | --- | --- | --- |
| HumanBaseTopGlobal [29] | ✓ | 25,689 | 77,807,094 | 0.117908 | Large & Dense |
| HuMAP [20] | ✓ | 15,433 | 35,052,604 | 0.147180 | Large & Dense |
| STRING [83] | ✓ | 18,480 | 11,019,492 | 0.032269 | Large |
| ConsensusPathDB [46] | ✓ | 17,735 | 10,611,416 | 0.033739 | Large |
| FunCoup [73] | ✓ | 17,892 | 10,037,478 | 0.031357 | Large |
| PCNet [36] | ✗ | 18,544 | 5,365,116 | 0.015603 | Large |
| BioGRID [82] | ✗ | 19,765 | 1,554,790 | 0.003980 | Medium |
| HumanNet [42] | ✓ | 18,591 | 2,250,780 | 0.006513 | Medium |
| HIPPIE [1] | ✓ | 19,338 | 1,542,044 | 0.004124 | Medium |
| ComPPIHumanInt [89] | ✓ | 17,015 | 699,620 | 0.002417 | Medium |
| OmniPath [85] | ✗ | 16,325 | 289,134 | 0.001085 | Small |
| ProteomeHD [50] | ✗ | 2,471 | 125,172 | 0.020509 | Small |
| HuRI [59] | ✗ | 8,100 | 103,188 | 0.001573 | Small |
| BioPlex [41] | ✗ | 8,108 | 71,004 | 0.001080 | Small |
| SIGNOR [70] | ✗ | 5,291 | 28,676 | 0.001025 | Small |

**Table S10:** Statistics for all datasets in OBNB (obnbdata-0.1.0)

| Label | Network | Num. tasks | Num. pos. avg. | Num. pos. std. | Num. pos. med. |
| --- | --- | --- | --- | --- | --- |
| DISEASES | BioGRID | 145 | 178.1 | 137.4 | 127.0 |
|  | BioPlex | 72 | 123.8 | 64.4 | 101.5 |
|  | ComPPIHumanInt | 145 | 174.6 | 134.5 | 125.0 |
|  | ConsensusPathDB | 144 | 177.4 | 137.5 | 126.0 |
|  | FunCoup | 145 | 177.1 | 135.1 | 127.0 |
|  | HIPPIE | 143 | 178.1 | 137.6 | 127.0 |
|  | HuMAP | 123 | 168.0 | 119.2 | 120.0 |
|  | HuRI | 50 | 130.3 | 56.7 | 112.5 |
|  | HumanBaseTopGlobal | 149 | 178.5 | 137.7 | 129.0 |
|  | HumanNet | 142 | 179.0 | 136.9 | 127.0 |
|  | OmniPath | 135 | 180.2 | 131.1 | 131.0 |
|  | PCNet | 143 | 171.8 | 130.6 | 122.0 |
|  | ProteomeHD | 15 | 76.9 | 22.4 | 70.0 |
|  | SIGNOR | 89 | 144.6 | 89.4 | 117.0 |
|  | STRING | 146 | 175.4 | 135.6 | 126.0 |
| DisGeNET | BioGRID | 305 | 208.3 | 143.1 | 159.0 |
|  | BioPlex | 189 | 138.6 | 71.4 | 111.0 |
|  | ComPPIHumanInt | 301 | 204.1 | 138.7 | 159.0 |
|  | ConsensusPathDB | 298 | 207.4 | 140.8 | 161.5 |
|  | FunCoup | 299 | 204.7 | 139.4 | 158.0 |
|  | HIPPIE | 306 | 208.1 | 142.9 | 159.5 |
|  | HuMAP | 279 | 194.3 | 126.7 | 155.0 |
|  | HuRI | 152 | 122.9 | 54.7 | 108.0 |
|  | HumanBaseTopGlobal | 287 | 219.7 | 145.7 | 173.0 |
|  | HumanNet | 302 | 204.2 | 140.3 | 158.5 |
|  | OmniPath | 298 | 199.6 | 136.0 | 153.5 |
|  | PCNet | 292 | 202.1 | 135.5 | 159.0 |
|  | ProteomeHD | 56 | 78.0 | 24.8 | 71.0 |
|  | SIGNOR | 219 | 147.3 | 81.9 | 124.0 |
|  | STRING | 296 | 208.0 | 140.6 | 162.0 |
| GOBP | BioGRID | 114 | 89.5 | 37.1 | 76.0 |
|  | BioPlex | 38 | 77.6 | 22.6 | 76.0 |
|  | ComPPIHumanInt | 104 | 91.8 | 37.0 | 77.5 |
|  | ConsensusPathDB | 112 | 90.1 | 37.0 | 76.5 |
|  | FunCoup | 114 | 87.8 | 36.7 | 74.0 |
|  | HIPPIE | 111 | 89.2 | 37.1 | 76.0 |
|  | HuMAP | 96 | 84.6 | 32.3 | 74.0 |
|  | HuRI | 27 | 69.9 | 16.0 | 65.0 |
|  | HumanBaseTopGlobal | 115 | 89.2 | 37.3 | 76.0 |
|  | HumanNet | 117 | 88.6 | 36.9 | 75.0 |
|  | OmniPath | 106 | 88.7 | 36.2 | 74.0 |
|  | PCNet | 105 | 89.0 | 36.0 | 77.0 |
|  | ProteomeHD | 5 | 80.4 | 22.6 | 70.0 |
|  | SIGNOR | 41 | 81.3 | 22.7 | 78.0 |
|  | STRING | 116 | 88.9 | 36.6 | 75.0 |
| GOCC | BioGRID | 71 | 96.0 | 36.3 | 87.0 |
|  | BioPlex | 35 | 77.0 | 20.9 | 71.0 |
|  | ComPPIHumanInt | 68 | 96.1 | 35.5 | 88.0 |
|  | ConsensusPathDB | 71 | 95.5 | 35.8 | 87.0 |
|  | FunCoup | 71 | 95.9 | 36.4 | 87.0 |
|  | HIPPIE | 69 | 97.3 | 35.8 | 89.0 |
|  | HuMAP | 62 | 91.8 | 33.4 | 82.0 |
|  | HuRI | 21 | 69.0 | 12.4 | 67.0 |
|  | HumanBaseTopGlobal | 71 | 96.9 | 36.3 | 88.0 |
|  | HumanNet | 70 | 95.8 | 35.6 | 88.0 |
|  | OmniPath | 67 | 94.0 | 34.6 | 84.0 |
|  | PCNet | 70 | 93.5 | 35.0 | 85.0 |
|  | ProteomeHD | 2 | 59.5 | 4.5 | 59.5 |
|  | SIGNOR | 26 | 66.4 | 11.3 | 67.5 |
|  | STRING | 69 | 96.1 | 35.6 | 88.0 |
| GOMF | BioGRID | 63 | 97.7 | 39.8 | 85.0 |
|  | BioPlex | 25 | 79.6 | 18.0 | 79.0 |
|  | ComPPIHumanInt | 62 | 98.1 | 39.7 | 86.0 |
|  | ConsensusPathDB | 63 | 98.2 | 39.5 | 85.0 |
|  | FunCoup | 64 | 97.5 | 39.1 | 83.5 |
|  | HIPPIE | 63 | 97.7 | 39.9 | 84.0 |
|  | HuMAP | 58 | 92.1 | 36.8 | 74.5 |
|  | HuRI | 22 | 74.8 | 12.8 | 73.5 |
|  | HumanBaseTopGlobal | 62 | 99.2 | 39.9 | 86.0 |
|  | HumanNet | 61 | 98.7 | 39.5 | 84.0 |
|  | OmniPath | 64 | 95.2 | 38.5 | 83.5 |
|  | PCNet | 58 | 97.4 | 38.6 | 86.0 |
|  | ProteomeHD | 2 | 59.5 | 4.5 | 59.5 |
|  | SIGNOR | 25 | 91.4 | 23.7 | 94.0 |
|  | STRING | 62 | 98.0 | 39.3 | 82.5 |

## Footnotes

1 The original paper uses mean aggregation, but we found that sum aggregation works better in our benchmark.

2 Uses the symmetric normalized adjacency matrix for propagation.

3 https://github.com/krishnanlab/obnbench/tree/main/conf

4 The area under the precision and recall curve.

5 The area under the receiver operator characteristic curve.

